# Age-related hyperexcitability in the mouse inferior colliculus: evidence from sound-evoked local field potentials

**DOI:** 10.64898/2026.09.04.749491

**Authors:** Dimitri L. Brunelle, Timothy J. Fawcett, Joseph P. Walton

## Abstract

The inferior colliculus (IC) is a major midbrain convergence site critical for processing complex sounds and undergoes fundamental changes with age-related hearing loss (ARHL). Local field potentials (LFPs) represent pre-synaptic integration of excitatory and inhibitory signals in local neural populations. Here, we assessed age-related changes in sound-evoked IC LFPs in the CBA/CaJ mouse model of ARHL across the tonotopic axis. We recorded from 495 sites across four age groups (young: 4-6 months; middle: 8-14 months; old: 24-25 months; “oldest old”: 27-31 months), examining LFP responses from dorsal (low-frequency), medial (mid-frequency), and ventral (high-frequency) IC regions. Analysis of the onset depolarization (N1 component) in response to broadband noise bursts revealed significant age-related hyperexcitability in medial and dorsal regions, with oldest old animals showing enhanced responses while ventral regions were unaffected. Amplitude-intensity functions demonstrated significant age × level interactions across all regions, with oldest old animals exhibiting level-dependent hyperexcitability most pronounced at suprathreshold intensities. Time-frequency analysis of LFP spectral content (30-100 Hz) revealed a crossover pattern across medial and dorsal regions, with oldest old animals showing reduced power at low-to-moderate stimulus levels but enhanced power at high stimulus levels. Ventral regions showed a low-level reduction without suprathreshold enhancement. These findings reveal paradoxical enhancement of neural activity in the aging IC, with the most pronounced changes in dorsal and medial regions rather than ventral regions most affected by peripheral hearing loss, suggesting that central hyperexcitability depends on residual afferent drive and implicating region-specific alterations in excitatory-inhibitory balance underlying central presbycusis.

## Introduction

Age-related hearing loss (ARHL), clinically termed presbycusis, is the gradual decline of hearing with age characterized by reduced hearing sensitivity and difficulty understanding speech in adverse listening situations (Gates and Mills, 2005; Humes and Dubno, 2010; Tremblay et al., 2003). ARHL is a major neurodegenerative disease impacting aging populations, affecting one-third of adults over the age of 65 (Lin et al., 2011b; Ohlemiller and Frisina, 2008; Schacht and Hawkins, 2005). Individuals with untreated hearing loss are more likely to experience social isolation and depression (Gates and Mills, 2005; Kalayam et al., 1995), both of which are risk factors for dementia and cognitive decline (Lin et al., 2011a). There are two major components to ARHL: loss in peripheral sensitivity (Lesicko and Llano, 2017; Lopez-Poveda and Barrios, 2013) and altered neural coding of sound in the central auditory system, decreasing the ability to localize sound and comprehend speech in noise (Auerbach and Gritton, 2022; Herrmann and Butler, 2021; Peelle and Wingfield, 2016; Resnik and Polley, 2021). Understanding this central component requires identifying where, and how, the aging brain transforms a degraded peripheral input into the neural representations that ultimately support sound perception.

The inferior colliculus (IC) is a major integrative gateway region of the brainstem (Oliver and Huerta, 1992) that has been shown to strongly influence central auditory processing (Oliver and Morest, 1984). The IC receives complex afferent input from both contralateral and ipsilateral nuclei (Frisina and Walton, 2001) and indirect ascending inputs from the auditory cortex (Oliver and Huerta, 1992; Schofield and Coomes, 2005), while also delivering efferent projections to the cochlear nucleus (Schofield and Cant, 1999). The projections from the lower auditory nuclei terminate at specific locations along the isofrequency planes of the IC (Brunso-Bechtold et al., 1981; Merzenich and Reid, 1974; Stiebler and Ehret, 1985), preserving tonotopic organization and allowing the IC to integrate excitatory and inhibitory input in a frequency-specific manner. Through this convergence, the IC plays an important role in coding virtually all dimensions of sound, including spectral content (De Martino et al., 2013; Egorova et al., 2001; Ehret and Merzenich, 1988; Stiebler and Ehret, 1985), temporal structure (Harris and Dubno, 2017; Schreiner and Langner, 1988; Schreiner and Winer, 2005; Walton et al., 1998), spatial location (Davis et al., 2003; Eramudugolla et al., 2008; McFadden and Willott, 1994; Semple et al., 1983), and species-specific vocalizations (Aitkin et al., 1994). Given this central role, age-related changes in the IC have been extensively studied in the context of neural presbycusis (Barsz et al., 2007; Leong et al., 2011; Ouda and Syka, 2012).

One underlying etiology of central presbycusis is a reduction in inhibitory neurotransmission within the central auditory system. Because inhibitory inputs play a critical role in shaping the frequency and temporal selectivity of IC neurons, their loss can produce broad changes in auditory processing, including reduced selectivity and exaggerated response magnitudes. Selective loss of GABAergic neurotransmission in the IC has been linked to auditory processing deficits in ARHL (Caspary et al., 1995; Caspary et al., 1990), and computational modeling of IC circuits indicates that simulated reductions in synaptic inhibition are sufficient to reproduce the altered, less selective response patterns observed in aged animals (Rabang et al., 2012). This loss of inhibitory tone is thought to act together with declining peripheral input to drive a compensatory increase in central gain, whereby central auditory structures upregulate their responsiveness as excitatory drive from the periphery weakens, producing disproportionately large responses to sound (Auerbach and Gritton, 2022; Rumschlag et al., 2022). Such central gain changes have been proposed to underlie paradoxical hyperactivity in the aging and hearing-impaired auditory system and may contribute to perceptual phenomena such as hyperacusis and tinnitus (Herrmann and Butler, 2021). Additionally, these neurochemical and circuit-level changes likely contribute to the temporal processing deficits associated with ARHL (Herrmann et al., 2017; Parthasarathy and Bartlett, 2011; Walton et al., 1998).

Extracellular neuronal signals are typically separated into spiking activity, which reflects the action potential output of a neuron, and slower potential fluctuations referred to as local field potentials (LFPs). While previous studies on sensory representations have focused primarily on spiking activity, the importance of field potentials in sensory processing has been increasingly recognized (Buzsáki et al., 2012; Einevoll et al., 2013). LFPs reflect synchronized pre-synaptic and dendritic activity across local neural populations, providing information about network-level processing that may not be apparent from single-unit or multi-unit spiking alone. Of particular interest in auditory processing are high-frequency oscillations in the 30-100 Hz band, which reflect local network synchrony and are especially sensitive to the balance between excitatory and inhibitory drive (Buzsáki and Wang, 2012). Because this excitatory-inhibitory balance is precisely what is thought to be disrupted by the loss of inhibitory tone described above, changes in 30-100 Hz spectral power offer a sensitive, circuit-level readout of central presbycusis. To localize age-related changes to the IC and its afferents, we therefore used LFPs as a proxy for pre-synaptic input to IC neurons.

Several studies have recorded local field potentials from the IC (Herrmann et al., 2017; Parthasarathy et al., 2019; Patel et al., 2012; Wang et al., 2002), and a subset of these have used LFPs specifically to investigate age-related changes in pre-synaptic input within the IC (Bartlett et al., 2024; Herrmann et al., 2017; Parthasarathy et al., 2019; Syka, 2020; Xiong et al., 2017). However, these studies have generally relied on binary young-versus-aged comparisons in rats, leaving open how LFP signatures evolve across the full lifespan and whether age effects differ across the tonotopic axis of the IC. In the present study, we examined LFP responses in the IC of CBA/CaJ mice across four age groups spanning the lifespan, sampled from dorsal (low-frequency), medial (mid-frequency), and ventral (high-frequency) IC. Using time-frequency analysis, we characterized the spectro-temporal properties of sound-evoked LFPs across these age groups, with particular focus on spectral activity in the 30-100 Hz range. Our findings reveal a paradoxical, region-specific enhancement of neural activity with advancing age, providing new insight into how central gain and disinhibition jointly reshape auditory midbrain processing across the lifespan.

## Experimental Procedures

### Subjects

CBA/CaJ mice were selected for this study because they experience decreased auditory function with age in a similar pattern to humans, making them a good model for the study of ARHL (Brunelle et al., 2025; Erway et al., 1993; Ohlemiller, 2006; Park et al., 2024; Willott et al., 1991). We used a total of 41 CBA/CaJ mice of both sexes spanning 495 units (female; N = 19, male; N = 22; see Supplemental Table S1.1 for counts across age groups). Four different age groups were used: young (4-6 months old; N = 7; 82 units), middle (8-14 months old; N = 7; 85 units), old (24-25 months old; N = 20; 243 units), and “oldest old” (27-31 months; N = 7; 85 units). A larger cohort of old animals was included to ensure adequate statistical power at the age range most commonly characterized as “aged” in the auditory aging literature. The oldest old cohort was necessarily smaller, as relatively few CBA/CaJ animals survive to 27-31 months in colony conditions. Regional sampling was not uniform across age groups; the number of animals contributing recordings to each region ranged from 4 to 20 (Supplemental Tables S1.2 and S1.3). Mice were raised in our in-house colony and were derived from breeders originally purchased from the Jackson Laboratory. Breeding pairs were reconstituted every 6-8 months. Mice were housed 3-4 per cage with litter-mates, in Seal-Safe Plus GM500 cages (36.9 × 15.6 × 13.2 cm) connected to a Box110SS Techniplast ventilated cage rack (West Chester, PA). The vivarium ambient noise levels across the frequency spectrum were as follows: 34.5 dB SPL at 2 kHz, 29 dB SPL at 4 kHz, 16 dB SPL at 8 kHz, 2.7 dB SPL at 16 kHz, and 0.4 dB SPL at 20 kHz. These measurements were taken from inside a cage located in the housing rack using a Quest Electronics Model 1800 sound level meter with the Model OB-300 1/3 octave filter set and a GRAS 40AE ½” Prepolarized Free-field microphone. The sound level meter was calibrated prior to recording with a Quest Electronics CA-22 Pistonphone Calibrator. The housing cages were maintained at a constant temperature (24 ± 1°C) and humidity (65 ± 3%), using a 12-hour light-dark cycle (7 A.M. to 7 P.M.) with water and food pellets available *ad libitum*. Electrophysiological testing occurred during the light cycle. Animals were closely monitored throughout the study period. All research was conducted following the University of South Florida Institutional Animal Care and Use Committee (IACUC) guidelines and was consistent with US Federal and NIH guidelines under IACUC protocol IS00014319, which was approved for renewal on 08/19/2025.

### Surgical preparation

Surgical procedures were conducted similarly to the reported methodology described in detail in previous studies from our laboratory (Barsz et al., 2007; Brecht et al., 2017; Brecht et al., 2022; Leong et al., 2011; Scott et al., 2017), details of which are reproduced here for clarity. Mice were initially anesthetized with an intraperitoneal injection containing 100 mg/kg ketamine and 10mg/kg xylazine. A test of the pedal reflex was used to assess the level of anesthesia. Once the mouse was completely anesthetized, the top of the mouse’s head and neck was shaved of fur. The bare skin was then cleaned using germicidal scrub, followed by 70% alcohol and iodine to ensure that the surgical area remained uncontaminated. Surgical scissors were then used to remove the bare skin, exposing the skull. The pericranium was scraped off using a scalpel, and 2% lidocaine was applied to the exposed tissue to further numb the area. A small hole was drilled and a ground electrode was placed on the contralateral parietal bone. A brass head post was then secured at the sagittal suture at bregma with Vetbond. Dental acrylic was spread over the remaining exposed skull and around the brass post to further secure the head post and ground electrode. Mice were given a recovery period of 24-48 hours before beginning the experimental sessions.

### Recording procedure

Prior to recording, chlorprothixene (Taractan, 5-12 µg/g, i.m.) was administered to induce sedation and prevent movement. Animals were secured in a custom stereotaxic frame (Newport-Klinger) within a heated (34 °C) sound attenuated booth lined with sound-absorbing foam (Sonex). The left IC was located stereotaxically (Paxinos and Franklin, 2007) and exposed via a small (<1 mm) craniotomy. Multi-unit extracellular activity was recorded using vertically oriented, single shank silicon acute penetrating 16-channel electrodes with an impedance ranging from 1.2 to 2.1 MΩ (Type-A, 3mm × 100 µm; 150 µm spacing, NeuroNexus Technologies). The electrode was positioned stereotaxically over the IC with reference to the lambda landmark and advanced dorsoventrally into the IC by a micro positioner (Newport-Klinger PMC 100). At least 4 electrode passes were made to map the extent of the IC and to identify the central nucleus. The output from the electrode was attached to a low noise (5-6 µV noise floor) preamplifier (RA16), having an operating range of ±7 mV. Neural events were acquired and visualized in real-time using the OpenEx software platform from Tucker-Davis Technologies (TDT, Inc., Alachua, FL) and a custom designed MATLAB® (The MathWorks, Inc., Natick, MA) graphical user interface. Neural recordings from each channel were then filtered (300-3000 Hz), amplified, and sampled at 25 kHz in a 1.25 ms time window subsequent to an event crossing a voltage discriminator. A spike triggering threshold of 4:1 signal to noise ratio (SNR) was automatically set for all channels. The search signal used to set the spike triggering thresholds was a 50 ms broadband noise (2-64 kHz) stimulus presented at 60 dB SPL at a rate of 5/s (up to 80 dB SPL depending on the mouse’s degree of hearing loss). Each penetration typically yielded 8-12 active channels. Recording sessions lasted an average of 6-8 hours, and if at any time a mouse showed signs of discomfort, such as excessive movement, it was removed from the recording apparatus and testing was halted.

### Stimulus generation and presentation

Noise and tone bursts were generated digitally (Real-time Processor Visual Design Studio (RVPds), TDT) using a System-3 processor and D/A converter (TDT RX6) with a 200 kHz sampling rate. The signals were routed to an electrostatic speaker (TDT ES1) with a flat frequency response from 4 to 110 kHz. This speaker was placed at 60° azimuth, contralateral to the recording site. Harmonic distortions were measured with a Dynamic Signal Analyzer (HP 35665A) and were at least 60 dB below the primary signal. The distance between the speaker and the pinna was fixed at 22.5 cm and calibrated using a B&K 2610 amplifier and a ¼” microphone placed at the location of the pinna. Excitatory frequency response areas (eFRAs) from all active channels were acquired simultaneously using 25 ms (5 ms rise/fall) tone burst stimuli presented from 0-80 dB SPL (5 dB steps) and from 2-64 kHz (log-spaced) for a total of 2125 frequency and intensity combinations that were presented pseudo-randomly five times at a rate of 10/s. Rate-level intensity functions were acquired in response to 50 ms broadband noise bursts (2-64 kHz, 0-80 dB SPL, 10 dB steps).

### Post-processing, signal and data analysis

#### Signal processing and eFRA/LFP extraction

Local field potentials were extracted from the recorded signals by band-pass filtering (2-300 Hz) and down sampling to 1 kHz. The N1 component, representing the primary depolarization response to stimulus onset, was identified as the largest negative deflection occurring within 8-12 ms post-stimulus. An automated peak detection algorithm was employed to ensure consistent identification across trials. N1 amplitude was measured as the absolute difference between baseline (pre-stimulus mean) and peak negative deflection. eFRAs were analyzed using a custom MATLAB program that classified tuning using a method similar to that used to classify neurons in the primary auditory cortex (Sutter, 2000). The frequency at which driven activity was responsive at the lowest intensity (threshold) was classified as the characteristic frequency, and the point in the receptive field which elicited maximal driven activity was categorized as the best frequency (BF). A custom MATLAB program was used to calculate the edges of each channel’s eFRA, which was then verified via visual inspection to ensure no non-driven activity was included in the calculation. The edges of the eFRA were defined as the activity levels that were equal to or greater than the background spike rate and at least 15% of the maximum spike rate. The data was grouped by electrode location into dorsal (channels 1-5), medial (channels 6-11), and ventral (channels 12-16) IC.

#### Time-frequency analysis

For time-frequency analysis, we employed a continuous Morlet wavelet transform with a constant ratio of f/Δf = 6.28 (Tallon-Baudry et al., 1996). The continuous wavelet transform was calculated using the formula:

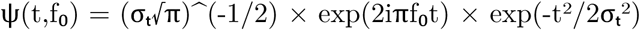

where f₀ is the center frequency, σₜ = 1/(2πσf) defines the temporal window, and σf = f₀/6.28 ensures constant relative bandwidth across frequencies.

#### Spectral power calculation

Wavelet convolution was performed on each trial, with time-frequency representations calculated for the 0-100 ms time interval and the 2-200 Hz frequency range at 1 ms time bins. Power at each time-frequency point was calculated as the squared magnitude of the complex wavelet coefficients. To analyze frequency-specific neural activity, power was averaged across the 30-100 Hz frequency band for each time point. We focused our analysis on the early response period (0-50 ms post-stimulus onset), which encompasses the N1 component of the LFP. This approach provides time-frequency estimates adapted to each frequency, though power estimates at the lower boundary of the band (30-40 Hz) reflect approximately 1-2 cycles and should be interpreted with this temporal constraint in mind.

#### Statistical analysis

Statistical analyses were performed using *R*, including the *data.table* and *dplyr* packages for data transformation. Multiple recording passes were typically obtained from the IC of each animal to sample across the tonotopic axis. Penetrations with the highest functional yield were selected to maximize spatial coverage and IC location. Only penetrations with stable recordings across the full depth of the IC and clear tonotopic gradients were included in the dataset. Given this hierarchical data structure (multiple recording sites nested within animals), recording sites served as the unit of analysis. For analysis of amplitude-intensity functions for both N1 amplitude and 30-100 Hz spectral power, we employed a two-way repeated measures ANOVA with age group as a between-subjects factor and stimulus level as a within-subjects factor, conducted separately for each frequency region (*aov()* function and *Error()* term in base *R*). To further characterize age effects at individual stimulus levels and for comparisons at single intensity levels (e.g., 80 dB SPL), we employed the Kruskal-Wallis test followed by post hoc pairwise Wilcoxon rank-sum tests with Holm-Bonferroni correction for multiple comparisons (*rstatix* package). This non-parametric approach was chosen due to heterogeneity of variance across age groups. Kruskal-Wallis results are reported as H(df), where H is the test statistic and significance is evaluated against a chi-square distribution with k -1 degrees of freedom. Alpha was set at 0.05 for all statistical tests. ANOVA model outputs were extracted and formatted using the *broom* package.

## Results

### IC LFPs are shaped by tonotopic organization and age-related changes

First, we investigated how IC tonotopic organization contributed to LFP morphology. Overall, minimum threshold was positively correlated with BF across all units (r = 0.184, p < 0.001) (Fig. 1A). This relationship was maintained in middle-aged (r = 0.376, p < 0.001) and old animals (r = 0.333, p < 0.001), but was absent in young (r = 0.206, p = 0.064) and oldest old animals (r = -0.098, p = 0.371) (Fig. 1A). Additionally, minimum threshold increased significantly with age (H(3) = 134.37, p < 0.001), from 28.9 ± 18.3 dB SPL in young and 25.4 ± 16.0 dB SPL in middle-aged animals to 36.5 ± 14.8 dB SPL in old and 54.0 ± 10.6 dB SPL in oldest old animals. Young and middle-aged groups did not differ in minimum threshold (p = 0.293); all other comparisons were significant (all p < 0.001). BF showed a strong tonotopic gradient across regions (Kruskal-Wallis: H(2) = 75.1, p < 0.001), with the ventral region having the highest BF (mean = 30.5 kHz), followed by the medial (mean = 19.9 kHz) and dorsal region (mean = 13.6 kHz) (Fig. 1B). This tonotopic organization was preserved across all age groups (Young, Middle, Old; all p < 0.001), though the oldest old group showed reduced regional differentiation (p = 0.091). While BF did not differ significantly across age groups overall (H(3) = 5.321, p = 0.150), channel-wise LFPs reveal age-related changes in morphology based on the respective unit’s characteristic frequency (Fig. 1C). Representative LFPs from a single representative mouse from each of the four age groups demonstrate an effect of frequency region on LFP morphology, with clear differences in waveform morphology characteristics across the tonotopic organization of the inferior colliculus. Fig. 1D shows the same units in Fig. 1C averaged across IC tonotopic subdivision, highlighting distinctions on N1 component characteristics in magnitude and latency between age groups. These findings are consistent with the expected high-frequency hearing loss characteristic of presbycusis in the IC.

**Fig. 1.**
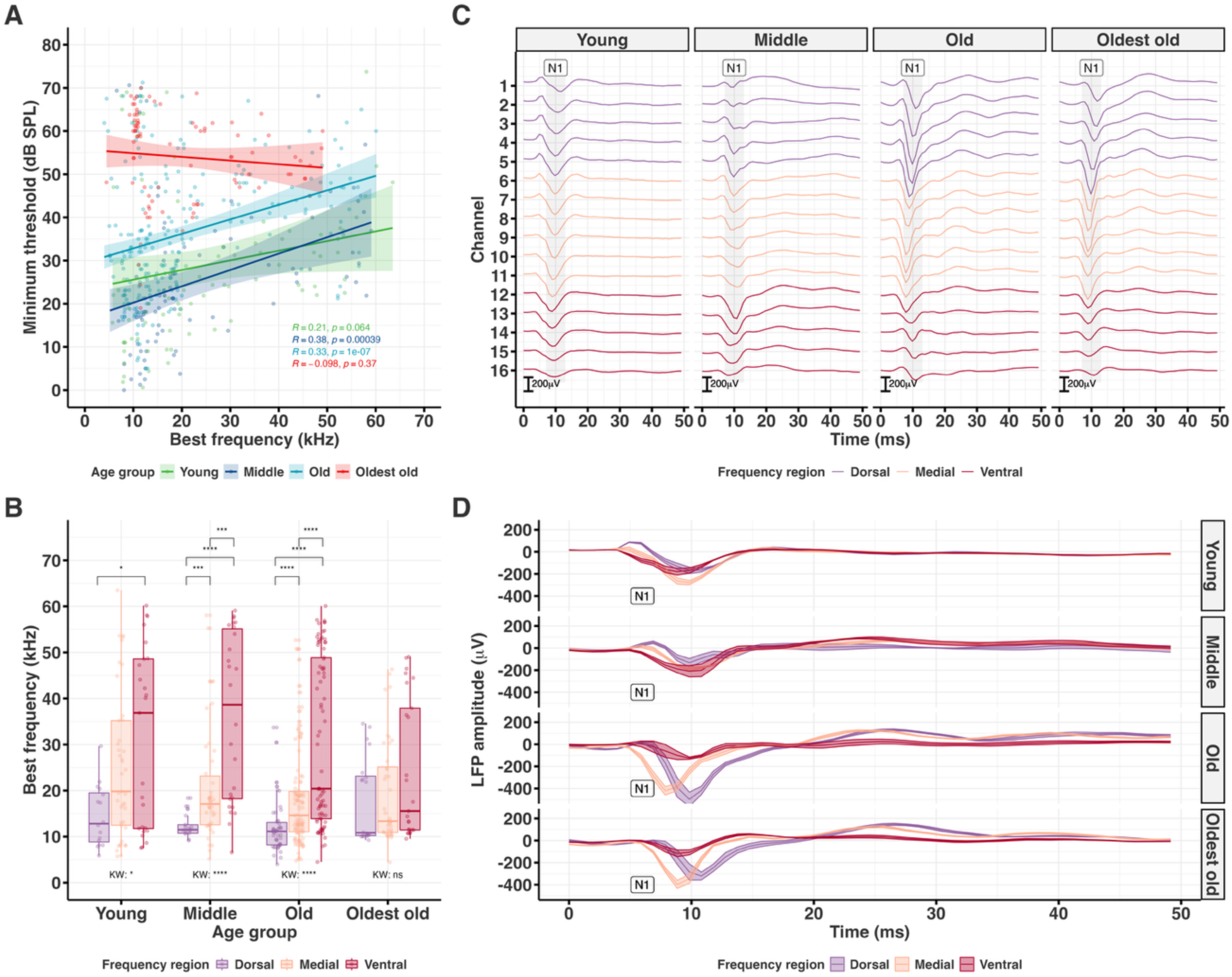
Frequency encoding characteristics and representative LFP waveforms in the aging inferior colliculus. **(A)** Relationship between minimum threshold and BF across all units, showing positive overall correlation (r = 0.184, p < 0.001***). Age group-wise correlations are shown in the figure inset. **(B)** BF distribution across anatomical regions and age groups. Strong tonotopic gradient is evident (Kruskal-Wallis H(2) = 75.1, p < 0.001***), with ventral regions showing highest BF (30.5 kHz), medial intermediate (19.9 kHz), and dorsal lowest (13.6 kHz). Within each age group, regional differences were tested using Kruskal-Wallis tests, with pairwise comparisons via Wilcoxon Rank Sum Test (Holm-Bonferroni correction). Brackets denote significance level of pairwise comparisons. Tonotopic organization is preserved in young, middle, and old groups (all p < 0.001***) but reduced in oldest old (p = 0.091, ns). **(C)** Representative channel-wise LFPs from a single representative mouse from each of the four age groups. The waveform component of interest is highlighted by gray windows indicating N1 (onset depolarization). Channel positions correspond to anatomical regions: channels 1-5 (dorsal), 6-11 (medial), 12-16 (ventral). **(D)** Grand averaged LFPs across anatomical regions for the units in **C** with N1 labeled. An effect of anatomical region on LFP morphology is observable. Shaded region represents SEM. See Supplemental Tables S2.1-S2.13 for all statistics pertaining to this figure.

### LFP N1 component is amplified in aged mice

Grand-average LFPs at 80 dB SPL across age groups and frequency regions are displayed in Fig. 2. The N1 component, occurring approximately 8-12 ms post-stimulus onset, represented the primary depolarization response and served as our primary measure of pre-synaptic activity and neural excitability. Qualitative examination of grand-averaged waveforms demonstrated pronounced enhancement of the N1 component with advancing age across the four age groups, particularly evident in the oldest old group (Fig. 2A). This enhancement was most pronounced in the dorsal and medial regions, where oldest old animals exhibited significantly larger negative deflections compared to younger cohorts.

**Fig. 2.**
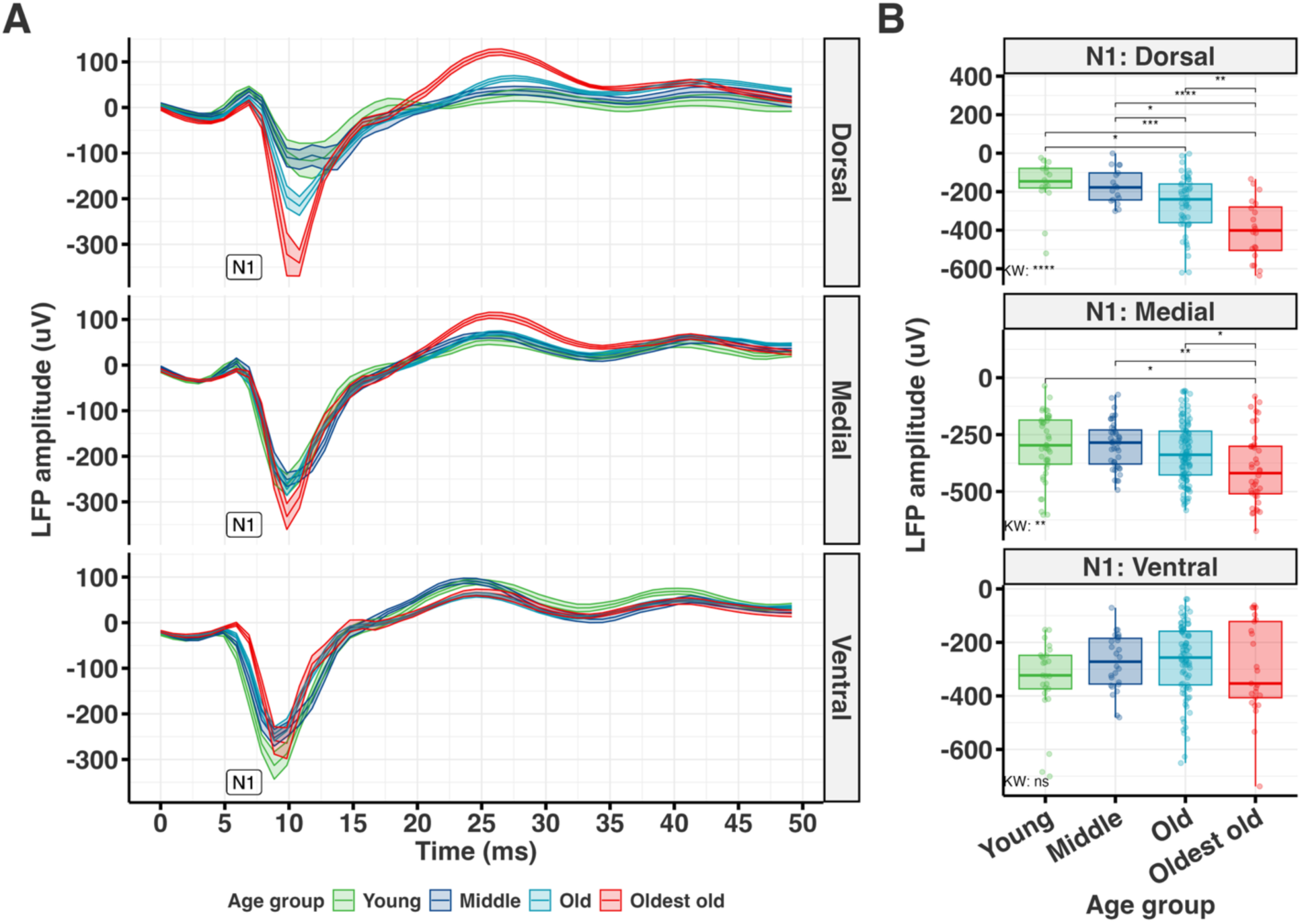
Grand-average LFP waveforms and amplitudes across anatomical regions. **(A)** Grand-average LFPs at 80 dB SPL across age groups and anatomical regions. The primary component of interest is labeled N1 (onset depolarization). An effect of age on LFP morphology is observable, with the oldest old group exhibiting a stronger depolarization profile relative to other age groups in the dorsal and medial regions. Shaded areas represent SEM. **(B)** Boxplots of N1 amplitude extracted from each unit’s LFP at 80 dB SPL across anatomical regions. Main effect of age was determined by the Kruskal-Wallis test (ventral: H(3) = 4.035, p = 0.258, ns; medial: H(3) = 14.251, p = 0.003**; dorsal: H(3) = 26.122, p < 0.001***). Significant age effects on N1 amplitude are evident in medial and dorsal regions, with oldest old animals showing greater depolarization consistent with age-related disinhibition in the midbrain. Pairwise comparisons were determined via Wilcoxon Rank Sum Test with Holm-Bonferroni correction. Brackets denote significance levels. See Supplemental Tables S3.1-S3.4 for all statistics pertaining to this figure.

Statistical analysis of N1 amplitudes at 80 dB SPL revealed significant age-related differences that varied by frequency region (Fig. 2B). In the ventral region, N1 amplitudes did not differ significantly between age groups (H(3) = 4.035, p = 0.258). However, both medial and dorsal regions demonstrated highly significant main effects of age (medial: H(3) = 14.251, p = 0.003; dorsal: H(3) = 26.122, p < 0.001), with the oldest old group showing substantially larger N1 amplitudes. Post-hoc pairwise comparisons revealed significant differences in the medial region between oldest old and young (p = 0.029), middle (p = 0.003), and old groups (p = 0.029). In the dorsal region, the pattern was even more pronounced, with oldest old differing significantly from young (p < 0.001), middle (p < 0.001), and old groups (p = 0.007). These differences represent significant increases in pre-synaptic depolarization, indicating fundamental changes in the balance of excitatory and inhibitory inputs to inferior colliculus neurons in advanced aging.

### LFP amplitude scales with stimulation level

Analysis of LFP responses across stimulus intensities (0-80 dB SPL) revealed significant differences in amplitude-intensity functions between age groups and anatomical regions (Fig. 3). While all age groups demonstrated the expected monotonic increase in absolute N1 amplitude with increasing stimulus level, the oldest old group exhibited distinctively steeper amplitude-intensity slopes, particularly in medial and dorsal regions. Two-way repeated measures ANOVA revealed significant main effects across all regions. In ventral and medial regions, both age (ventral: F(3, 154) = 8.576, p < 0.001; medial: F(3, 230) = 6.051, p < 0.001) and level (ventral: F(8, 1232) = 337.028, medial: F(8, 1840) = 627.033, both p < 0.001) were significant, along with significant Age × Level interactions (ventral: F(24, 1232) = 8.326, p < 0.001; medial: F(24, 1840) = 25.134, p < 0.001). In the dorsal region, while the main effect of age was not significant (F(3, 99) = 1.314, p = 0.274), the main effect of level was highly significant (F(8, 792) = 188.196, p < 0.001), as was the Age × Level interaction (F(24, 792) = 12.404, p < 0.001), indicating that age effects in this region are level-dependent.

**Fig. 3.**
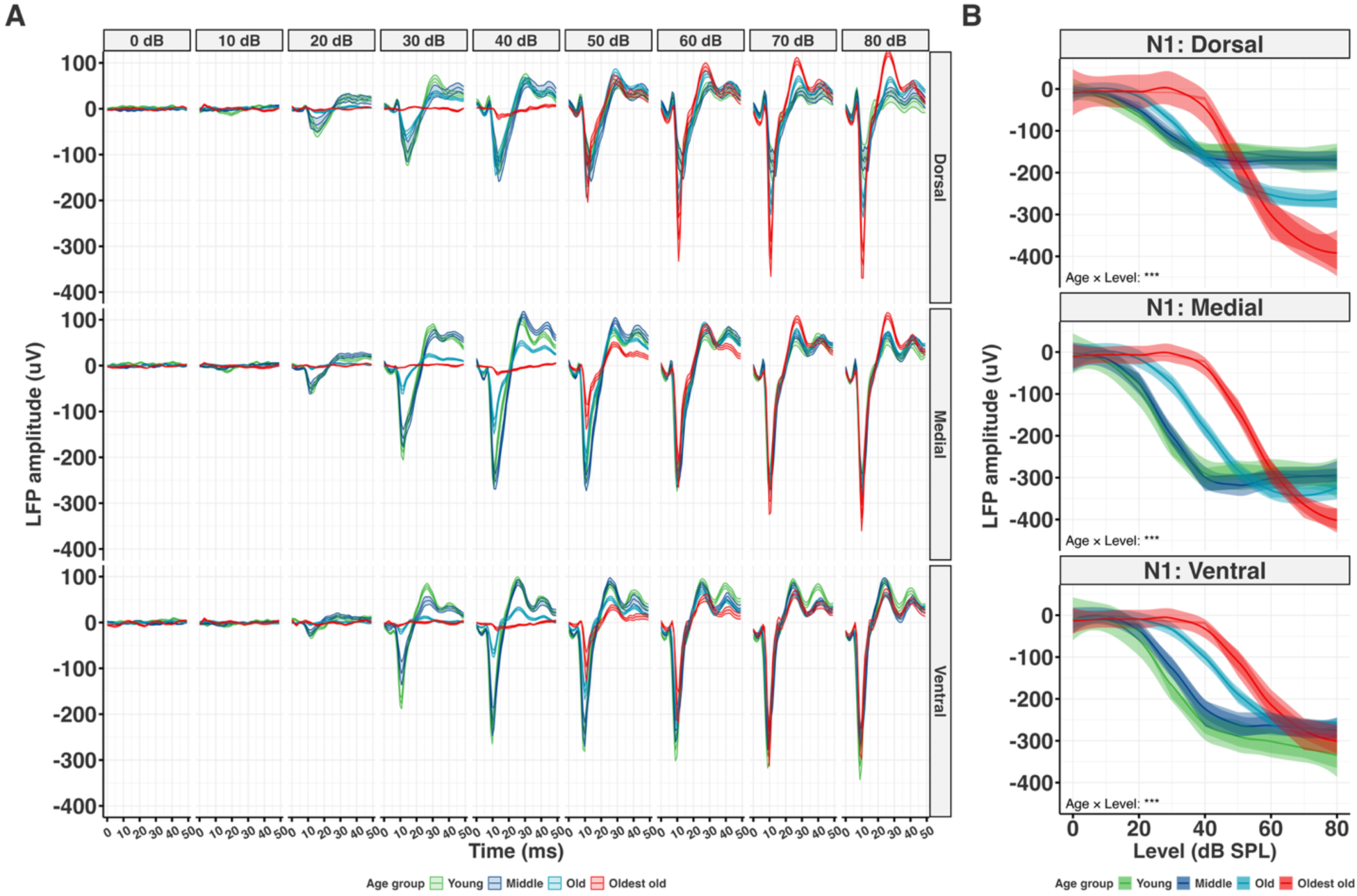
Effect of stimulus level on average LFP waveforms and amplitudes across age groups and anatomical regions. **(A)** Grand-average LFP waveforms across stimulus levels (0-80 dB SPL) showing progression of response magnitude with increasing intensity. Responses are organized by anatomical region (dorsal, medial, ventral) and age group. Shaded regions represent SEM. **(B)** N1 amplitude-intensity functions for broadband noise stimulation across anatomical regions. Two-way repeated measures ANOVA revealed significant Age × Level interactions in all regions (ventral: F(24, 1232) = 8.326, p < 0.001***; medial: F(24, 1840) = 25.134, p < 0.001***; dorsal: F(24, 792) = 12.404, p < 0.001***). The dorsal region shows the most dramatic age effect, with oldest old animals exhibiting significantly different responses across most stimulus levels tested (all p < 0.05 except 50 dB SPL, omnibus Kruskal-Wallis), indicating level-dependent hyperexcitability. Mean ± SEM are shown. See Supplemental Tables S4.1-S4.4 for all statistics pertaining to this figure.

Notably, the direction of age-related differences reversed across stimulus intensity, revealing a crossover pattern that differed in prominence by region. In ventral regions, oldest old animals showed significantly reduced N1 amplitudes relative to all younger groups from 30-50 dB SPL (all p ≤ 0.023; e.g., -261 µV in young vs. -19 µV in oldest old at 40 dB); at 20 dB SPL the reduction reached significance only against young animals (p < 0.001). At high levels (60-80 dB SPL), no ventral comparison survived correction, and oldest old amplitudes approached those of other groups without surpassing them. Medial regions showed a comparable pattern, with oldest old amplitudes significantly reduced relative to all younger groups from 30-50 dB SPL (all p < 0.001) and relative to young and middle-aged animals at 10-20 dB SPL. However, the pattern reversed at 80 dB SPL: oldest old animals exhibited the largest N1 amplitudes (-399 ± 156 µV vs. young -300 ± 145 µV), differing significantly from young (p = 0.029), middle (p = 0.003), and old (p = 0.029) groups. Dorsal regions showed the most pronounced crossover, with oldest old significantly reduced relative to all younger groups at 30-40 dB SPL (all p < 0.001), a non-significant transition at 50-60 dB SPL, and significant enhancement at 70-80 dB SPL, reaching the largest responses of any group (-397 ± 154 µV vs. young -164 ± 133 µV at 80 dB SPL; vs. young p < 0.001, vs. middle p < 0.001, vs. old p = 0.007). Post-hoc comparisons confirmed that oldest old animals differed significantly from all three younger groups at 80 dB SPL in both dorsal and medial regions (all p < 0.05, Holm-Bonferroni correction).

### LFP high-frequency spectral power increases with age in a region- and level-dependent manner

Time-frequency analysis was computed using Morlet wavelet transforms (f/Δf = 6.28) which extracted spectral content from LFP responses with optimal time-frequency resolution for neural signals (Fig. 4A). The analysis focused on high-frequency oscillations in the 30-100 Hz frequency band during the first 50 ms post-stimulus, capturing the onset-evoked spectral response that reflects local network synchrony and the balance between excitatory and inhibitory processes. Spectrograms revealed robust time-frequency representations across all age groups and stimulus conditions (Fig. 4B). The high-frequency power was maximal during the first 20 ms post-stimulus, coinciding with the N1 component, and showed clear intensity-dependent modulation (Fig. 4C). Visual inspection of time-frequency representations suggests enhanced 30-100 Hz power in oldest old animals, particularly in medial and dorsal regions of the IC.

**Fig. 4.**
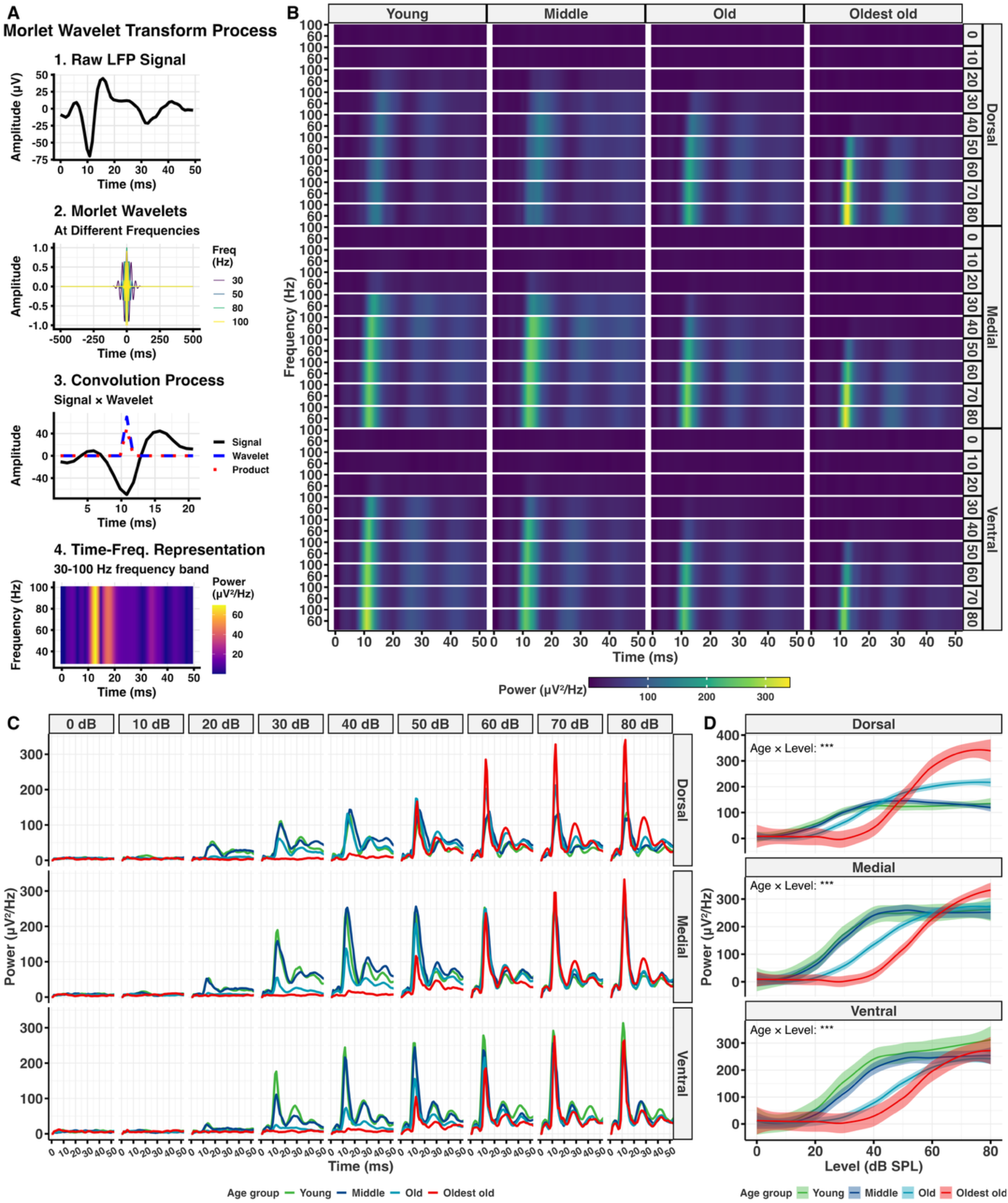
Time-frequency analysis reveals enhanced high-frequency neural synchronization in the aging inferior colliculus. **(A)** Morlet wavelet transform methodology showing the four-step process: (1) raw LFP signal, (2) Morlet wavelets at different frequencies (30, 50, 80, 100 Hz), (3) convolution process demonstrating signal × wavelet interaction, and (4) resulting time-frequency representation for the 30-100 Hz band. **(B)** Time-frequency spectrograms showing power in the 30-100 Hz range across age groups and stimulus levels (0-80 dB SPL) for all units from each anatomical region. Color scale represents power (µV²/Hz). Note the regional and age-dependent variations in 30-100 Hz power patterns. **(C)** Time course of averaged 30-100 Hz band power across stimulus levels and age groups for dorsal, medial, and ventral anatomical regions. Regional differences in age effects are evident, with complex level-dependent patterns. **(D)** Intensity functions for peak 30-100 Hz power across anatomical regions. Two-way repeated measures ANOVA revealed significant Age × Level interactions in all regions (ventral: F(18, 924) = 7.988, p < 0.001***; medial: F(18, 1380) = 31.658, p < 0.001***; dorsal: F(18, 594) = 16.266, p < 0.001***). In ventral regions, age effects are confined to low-to-moderate levels (20-50 dB SPL) with oldest old showing reduced 30-100 Hz power and no suprathreshold enhancement. Medial regions show the same low-level reduction followed by a crossover to enhanced power at 80 dB SPL. In dorsal regions, the crossover occurs at lower intensities: age effects are significant at 20-40 dB SPL (with oldest old showing reduced power) and at 60-80 dB SPL (with oldest old showing enhanced power), separated by a non-significant transition at 50 dB SPL. Mean ± SEM are shown. See Supplemental Tables S5.1-S5.4 for all statistics pertaining to this figure.

Quantitative analysis of peak high-frequency power (30-100 Hz) revealed significant age-related increases that paralleled the N1 amplitude findings across frequency regions and age (Fig. 4D). Ventral regions showed significant main effects of age (F(3, 154) = 8.248, p < 0.001), level (F(6, 924) = 262.214, p < 0.001), and their interaction (F(18, 924) = 7.988, p < 0.001). Medial regions showed significant main effects of age (F(3, 230) = 5.637, p < 0.001) and level (F(6, 1380) = 441.000, p < 0.001), together with a markedly larger Age × Level interaction than in ventral regions (F(18, 1380) = 31.658, p < 0.001), indicating a more pronounced level dependence of the age effect. Dorsal regions showed a pattern similar to N1 amplitude, with no main effect of age (F(3, 99) = 1.273, p = 0.288) but a significant main effect of level (F(6, 594) = 143.674, p < 0.001) and a highly significant Age × Level interaction (F(18, 594) = 16.266, p < 0.001). Post-hoc analyses revealed that age effects on 30-100 Hz power were highly level-dependent, with all three regions showing significantly reduced power in oldest old animals at low-to-moderate stimulus levels, and medial and dorsal regions additionally showing a crossover to enhanced power at high levels. In ventral and medial regions, significant reductions in 30-100 Hz power were evident at 20-50 dB SPL (all omnibus p < 0.001), with oldest old animals showing the lowest power values. At higher stimulus levels (60-70 dB SPL), these differences diminished in both regions, though medial regions showed a return to significance at 80 dB SPL (p = 0.003), where oldest old animals exhibited greater power than younger groups. In dorsal regions, the same crossover pattern was evident but shifted along the intensity axis. Significant reductions in oldest old animals occurred at 20-40 dB SPL (all p < 0.001), followed by a non-significant transition at 50 dB SPL (p = 0.111), and significant enhancement at 60-80 dB SPL (all p < 0.05), with oldest old animals exhibiting the largest responses. Notably, this dorsal crossover occurred one intensity step below the corresponding crossover in N1 amplitude (60 vs. 70 dB SPL; cf. Fig. 3), indicating that the transition to suprathreshold enhancement is detectable in spectral power slightly earlier along the intensity axis than in response magnitude. This shared low-level reduction across all three regions, together with a crossover to suprathreshold enhancement confined to dorsal (60-80 dB SPL) and medial (80 dB SPL) regions suggests a common alteration in intensity coding whose suprathreshold expression depends on residual peripheral drive.

## Discussion

The present study provides compelling evidence for increased hyperexcitability in sound-evoked IC LFPs in aged mice. Our findings reveal three primary changes associated with advanced aging: (1) significantly enhanced N1 depolarization responses in oldest old animals, particularly in medial and dorsal regions; (2) steeper amplitude-intensity functions indicating suprathreshold hyperexcitability; and (3) a level-dependent crossover in high-frequency spectral power (30-100 Hz), with oldest old animals showing reduced power at low-to-moderate levels but enhanced power at suprathreshold levels, suggesting altered intensity coding of local network synchrony. These results demonstrate a paradoxical enhancement of neural activity in the aging IC despite the presence of peripheral hearing loss, providing quantitative support for the disinhibition hypothesis of central presbycusis (Auerbach et al., 2014; Caspary et al., 1995; Ibrahim and Llano, 2019).

### Disinhibition, local circuit reorganization, and suprathreshold hyperexcitability

The striking enhancement of N1 amplitudes in oldest old animals provides direct evidence for altered excitatory-inhibitory balance in the aging IC. The N1 component reflects the summation of pre-synaptic depolarizations within local neural populations (Kajikawa and Schroeder, 2011), making it a sensitive measure of population-level excitability (Buzsáki et al., 2012; Einevoll et al., 2013). Prior LFP studies in the rat IC found onset-evoked LFP amplitudes to be larger in younger than aged animals at moderate stimulus intensities (Bartlett et al., 2024; Herrmann et al., 2017), a pattern we replicate at low-to-moderate levels across all three regions. However, our intensity-function approach additionally reveals a crossover to suprathreshold hyperexcitability that fixed-intensity paradigms have not previously characterized to our knowledge. These observations are consistent with the disinhibition hypothesis, which posits that age-related loss of GABAergic inhibition unmasks excitatory responses and reduces the gain control normally exerted across the dynamic range of stimulus intensities (Auerbach et al., 2014; Caspary et al., 2008; Caspary et al., 1995; Caspary et al., 1990; Ibrahim and Llano, 2019; Milbrandt et al., 1994; Ouda and Syka, 2012), supported pharmacologically by studies showing that augmenting GABAergic inhibition in aged IC neurons reverses age-related alterations in both frequency tuning and intensity coding (Brecht et al., 2017; Caspary et al., 2002). Loss of these inhibitory constraints produces broader, less selective responses with increased susceptibility to excitatory drive, which is consistent with single-unit studies showing decreased frequency selectivity and altered temporal processing in aged IC neurons (Leong et al., 2011; Palombi et al., 2001; Parthasarathy et al., 2019; Walton et al., 1998). Once threshold is exceeded, aged IC circuits amplify responses disproportionately as reflected in a report by Xiong et al. (2017), which demonstrated an association between IC spiking hyperexcitability and age-related cochlear hearing loss.

### Regional specificity of age-related changes

The age-related hyperexcitability showed clear regional specificity, with effects most pronounced in dorsal and medial IC. The dorsal region exhibited the most dramatic changes, with a highly significant Age × Level interaction despite no significant main effect of age, indicating that hyperexcitability emerges specifically at suprathreshold intensities. Ventral regions showed markedly smaller functional changes despite presumably experiencing greater peripheral hearing loss and the most pronounced neurochemical vulnerability. Prior studies have documented the greatest reductions in GABA and glycine immunoreactivity in high-frequency regions of the aging IC (Caspary et al., 1995; Milbrandt et al., 1994). This dissociation between neurochemical vulnerability and functional hyperexcitability in ventral regions may reflect ceiling effects in inhibitory loss, compensatory upregulation of remaining inhibitory circuits, or the dominance of peripheral deafferentation that prevents centrally-driven differences from manifesting. Prior IC LFP aging studies have typically recorded from single sites without examining regional differences across the tonotopic axis (Bartlett et al., 2024; Herrmann et al., 2017; Xiong et al., 2017), leaving open whether population-level hyperexcitability is uniform or region-selective. Our multi-site approach suggests that it is decidedly region-selective.

The level-dependent crossover in N1 amplitude with oldest old animals showing reduced responses at low-to-moderate levels and enhanced responses at suprathreshold levels in medial and dorsal regions suggests that peripheral and central factors dominate different portions of the intensity function. At low-to-moderate levels, reduced peripheral drive secondary to cochlear aging (Kujawa and Liberman, 2009) suppresses N1 amplitude across all regions; at suprathreshold levels, central disinhibition may overcome this deficit in medial and dorsal regions, producing greatly enhanced responses in oldest old animals. In ventral regions, the peripheral deficit persists across the full intensity range, consistent with the greater cochlear damage expected in high-frequency regions and within the central gain framework, in which compensatory central upregulation only manifests with sufficient residual peripheral drive to recruit disinhibited circuits (Chambers et al., 2016; Parthasarathy et al., 2019; Rumschlag et al., 2022).

### High-frequency spectral content and neural synchronization

The parallel crossover in 30-100 Hz spectral power that mirrors the N1 amplitude pattern across medial and dorsal regions extends the disinhibition framework from response magnitude to local network synchrony. 30-100 Hz oscillations reflect the synchronization of local neural populations arising from the precise timing of inhibitory interneuron activity (Bartos et al., 2007; Buzsáki and Wang, 2012), making their enhancement at suprathreshold levels apparently paradoxical given documented inhibitory loss. This paradox may be resolved by considering the complex relationship between inhibitory function and network synchrony - while overall inhibitory strength may be reduced, the remaining inhibitory circuits may exhibit altered dynamics that promote coherent population activity under strong excitatory drive (Isaacson and Scanziani, 2011). A related observation was made by Herrmann et al. (2017), who found enhanced spike synchronization at low modulation rates (45 Hz) in aged rat IC despite similar LFP synchronization between age groups over a 225 ms window, suggesting that the aged IC amplifies the transformation from synaptic input to synchronized spiking output. Our finding of enhanced 30-100 Hz spectral power at suprathreshold levels may reflect an analogous process, with strong excitatory drive enabling remaining inhibitory circuits to generate more coherent synchronized population activity. Alternatively, reduced inhibitory precision rather than reduced overall strength may yield more variable but sporadically hypersynchronous population responses, producing increased spectral power at high excitatory drive levels without reflecting a stable increase in circuit synchrony. Critically, the 30-100 Hz crossover pattern closely tracked the N1 amplitude crossover across regions, including the absence of suprathreshold enhancement in ventral regions. Because both measures derive from the same onset transient, this correspondence is expected - establishing that spectral and amplitude readouts index a common shift in excitatory-inhibitory balance which are gated by peripheral input strength.

### Functional implications for auditory processing

Our findings contribute to the growing recognition of paradoxical hyperactivity in sensory systems affected by peripheral impairment (Herrmann and Butler, 2021), with the level-dependent crossover providing a mechanistic bridge between peripheral hearing loss and its perceptual consequences. The crossover likely predicts intensity-dependent perceptual effects. Reduced neural responses at near-threshold levels likely contribute to difficulty detecting and processing soft sounds, while disproportionately large responses at suprathreshold levels may underlie loudness recruitment and reduced tolerance for intense sounds, both common complaints in aging, as well as perceptual phenomena such as hyperacusis and tinnitus that have been linked to similar disinhibition mechanisms across forms of hearing loss (Herrmann and Butler, 2021; Llano et al., 2012). Increased but less selective responses may further contribute to difficulty understanding speech in background noise, consistent with behavioral signal-in-noise detection deficits across the lifespan in the CBA/CaJ mouse model (Anderson et al., 2012; Brunelle et al., 2025; Parthasarathy et al., 2019). Several considerations bear on the interpretation of these findings. Recording sites served as the unit of analysis and regional sampling was uneven across age groups. Recordings were also obtained in sedated rather than fully awake animals; while this avoids the response suppression associated with general anesthesia, age-related differences in sensitivity to sedation cannot be excluded. Whether the changes documented here represent adaptive homeostatic compensation for reduced peripheral input or an age-related maladaptive disruption of auditory processing precision remains an important open question (Ibrahim and Llano, 2019) - one that future studies combining behavioral and neural measures across the full intensity range may help resolve.

## Supporting information

Supplemental Statistical Analyses

## CRediT authorship contribution statement

**Dimitri L. Brunelle**: Writing - review & editing, Writing - original draft, Visualization, Validation, Software, Methodology, Investigation, Formal analysis, Data curation. **Timothy J. Fawcett**: Writing - review & editing, Validation, Software, Methodology, Formal analysis, Conceptualization. **Joseph P. Walton**: Writing - review & editing, Validation, Project administration, Supervision, Resources, Methodology, Funding acquisition, Conceptualization.

## Declaration of competing interest

The authors declare that they have no known competing financial interests or personal relationships that could have appeared to influence the work reported in this paper.

## Acknowledgements

We thank Drs. Michelle R. Kapolowicz and Erol J. Ozmeral in the Department of Communication Sciences and Disorders along with Dr. Robert D. Frisina in the Department of Medical Engineering at the University of South Florida for their valuable expertise and feedback on the study.

## Notes

### Competing Interest Statement

The authors have declared no competing interest.

## References

Aitkin, L., Tran, L., Syka, J., 1994. The responses of neurons in subdivisions of the inferior colliculus of cats to tonal, noise and vocal stimuli. Exp Brain Res 98(1), 53–64.

Anderson, S., Parbery-Clark, A., White-Schwoch, T., Kraus, N., 2012. Aging affects neural precision of speech encoding. Journal of Neuroscience 32(41), 14156–14164.

Auerbach, B.D., Gritton, H.J., 2022. Hearing in Complex Environments: Auditory Gain Control, Attention, and Hearing Loss. Front Neurosci 16, 799787.

Auerbach, B.D., Rodrigues, P.V., Salvi, R.J., 2014. Central gain control in tinnitus and hyperacusis. Frontiers in neurology 5, 206.

Barsz, K., Wilson, W.W., Walton, J.P., 2007. Reorganization of receptive fields following hearing loss in inferior colliculus neurons. Neuroscience 147(2), 532–545.

Bartlett, E.L., Han, E.X., Parthasarathy, A., 2024. Neurometric amplitude modulation detection in the inferior colliculus of Young and Aged rats. Hearing research 447, 109028.

Bartos, M., Vida, I., Jonas, P., 2007. Synaptic mechanisms of synchronized gamma oscillations in inhibitory interneuron networks. Nature reviews neuroscience 8(1), 45–56.

Brecht, E.J., Barsz, K., Gross, B., Walton, J.P., 2017. Increasing GABA reverses age-related alterations in excitatory receptive fields and intensity coding of auditory midbrain neurons in aged mice. Neurobiology of aging 56, 87–99.

Brecht, E.J., Scott, L.L., Ding, B., Zhu, X., Walton, J.P., 2022. A BK channel-targeted peptide induces age-dependent improvement in behavioral and neural sound representation. Neurobiology of Aging 110, 61–72.

Brunelle, D.L., Park, C.R., Fawcett, T.J., Walton, J.P., 2025. Signal-in-noise detection across the lifespan in a mouse model of presbycusis. Hearing Research 455, 109153.

Brunso-Bechtold, J.K., Thompson, G.C., Masterton, R.B., 1981. HRP study of the organization of auditory afferents ascending to central nucleus of inferior colliculus in cat. J Comp Neurol 197(4), 705–722.

Buzsáki, G., Anastassiou, C.A., Koch, C., 2012. The origin of extracellular fields and currents— EEG, ECoG, LFP and spikes. Nature reviews neuroscience 13(6), 407–420.

Buzsáki, G., Wang, X.-J., 2012. Mechanisms of gamma oscillations. Annual review of neuroscience 35(1), 203–225.

Caspary, D.M., Ling, L., Turner, J.G., Hughes, L.F., 2008. Inhibitory neurotransmission, plasticity and aging in the mammalian central auditory system. Journal of Experimental Biology 211(11), 1781–1791.

Caspary, D.M., Milbrandt, J.C., Helfert, R.H., 1995. Central auditory aging: GABA changes in the inferior colliculus. Exp Gerontol 30(3-4), 349–360.

Caspary, D.M., Palombi, P.S., Hughes, L.F., 2002. GABAergic inputs shape responses to amplitude modulated stimuli in the inferior colliculus. Hearing research 168(1-2), 163–173.

Caspary, D.M., Raza, A., Lawhorn Armour, B.A., Pippin, J., Arneric, S.P., 1990. Immunocytochemical and neurochemical evidence for age-related loss of GABA in the inferior colliculus: implications for neural presbycusis. J Neurosci 10(7), 2363-2372.

Chambers, A.R., Resnik, J., Yuan, Y., Whitton, J.P., Edge, A.S., Liberman, M.C., Polley, D.B., 2016. Central gain restores auditory processing following near-complete cochlear denervation. Neuron 89(4), 867–879.

Davis, K.A., Ramachandran, R., May, B.J., 2003. Auditory processing of spectral cues for sound localization in the inferior colliculus. J Assoc Res Otolaryngol 4(2), 148–163.

De Martino, F., Moerel, M., van de Moortele, P.F., Ugurbil, K., Goebel, R., Yacoub, E., Formisano, E., 2013. Spatial organization of frequency preference and selectivity in the human inferior colliculus. Nat Commun 4(1), 1386.

Egorova, M., Ehret, G., Vartanian, I., Esser, K.H., 2001. Frequency response areas of neurons in the mouse inferior colliculus. I. Threshold and tuning characteristics. Exp Brain Res 140(2), 145–161.

Ehret, G., Merzenich, M.M., 1988. Complex sound analysis (frequency resolution, filtering and spectral integration) by single units of the inferior colliculus of the cat. Brain Res 472(2), 139–163.

Einevoll, G.T., Kayser, C., Logothetis, N.K., Panzeri, S., 2013. Modelling and analysis of local field potentials for studying the function of cortical circuits. Nat Rev Neurosci 14(11), 770–785.

Eramudugolla, R., McAnally, K.I., Martin, R.L., Irvine, D.R., Mattingley, J.B., 2008. The role of spatial location in auditory search. Hear Res 238(1-2), 139–146.

Erway, L.C., Willott, J.F., Archer, J.R., Harrison, D.E., 1993. Genetics of age-related hearing loss in mice: I. Inbred and F1 hybrid strains. Hearing research 65(1-2), 125–132.

Frisina, R.D., Walton, J.P., 2001. Neuroanatomy of the central auditory system, Handbook of Mouse Auditory Research. CRC Press, pp. 257–292.

Gates, G.A., Mills, J.H., 2005. Presbycusis. The lancet 366(9491), 1111–1120.

Harris, K.C., Dubno, J.R., 2017. Age-related deficits in auditory temporal processing: unique contributions of neural dyssynchrony and slowed neuronal processing. Neurobiology of aging 53, 150–158.

Herrmann, B., Butler, B.E., 2021. Hearing loss and brain plasticity: the hyperactivity phenomenon. Brain Struct Funct 226(7), 2019–2039.

Herrmann, B., Parthasarathy, A., Bartlett, E.L., 2017. Ageing affects dual encoding of periodicity and envelope shape in rat inferior colliculus neurons. Eur J Neurosci 45(2), 299–311.

Humes, L.E., Dubno, J.R., 2010. Factors Affecting Speech Understanding in Older Adults, in: Gordon-Salant, S., Frisina, R.D., Popper, A.N., Fay, R.R. (Eds.), The Aging Auditory System. Springer New York, New York, NY, pp. 211–257.

Ibrahim, B.A., Llano, D.A., 2019. Aging and central auditory disinhibition: is it a reflection of homeostatic downregulation or metabolic vulnerability? Brain Sciences 9(12), 351.

Isaacson, J.S., Scanziani, M., 2011. How inhibition shapes cortical activity. Neuron 72(2), 231–243.

Kajikawa, Y., Schroeder, C.E., 2011. How local is the local field potential? Neuron 72(5), 847–858.

Kalayam, B., Meyers, B.S., Kakuma, T., Alexopoulos, G.S., Young, R.C., Solomon, S., Shotland, R., Nambudiri, D., Goldsmith, D., 1995. Age at onset of geriatric depression and sensorineural hearing deficits. Biological Psychiatry 38(10), 649–658.

Kujawa, S.G., Liberman, M.C., 2009. Adding insult to injury: cochlear nerve degeneration after “temporary” noise-induced hearing loss. Journal of Neuroscience 29(45), 14077–14085.

Leong, U.C., Barsz, K., Allen, P.D., Walton, J.P., 2011. Neural correlates of age-related declines in frequency selectivity in the auditory midbrain. Neurobiol Aging 32(1), 168–178.

Lesicko, A.M., Llano, D.A., 2017. Impact of peripheral hearing loss on top-down auditory processing. Hear Res 343, 4–13.

Lin, F.R., Metter, E.J., O’Brien, R.J., Resnick, S.M., Zonderman, A.B., Ferrucci, L., 2011a. Hearing loss and incident dementia. Archives of neurology 68(2), 214–220.

Lin, F.R., Niparko, J.K., Ferrucci, L., 2011b. Hearing loss prevalence in the United States. Archives of internal medicine 171(20), 1851–1853.

Llano, D.A., Turner, J., Caspary, D.M., 2012. Diminished cortical inhibition in an aging mouse model of chronic tinnitus. Journal of Neuroscience 32(46), 16141–16148.

Lopez-Poveda, E.A., Barrios, P., 2013. Perception of stochastically undersampled sound waveforms: a model of auditory deafferentation. Front Neurosci 7, 124.

McFadden, S.L., Willott, J.F., 1994. Responses of inferior colliculus neurons in C57BL/6J mice with and without sensorineural hearing loss: effects of changing the azimuthal location of a continuous noise masker on responses to contralateral tones. Hear Res 78(2), 132–148.

Merzenich, M.M., Reid, M.D., 1974. Representation of the cochlea within the inferior colliculus of the cat. Brain research 77(3), 397–415.

Milbrandt, J.C., Albin, R.L., Caspary, D.M., 1994. Age-related decrease in GABAB receptor binding in the Fischer 344 rat i inferior colliculus. Neurobiology of aging 15(6), 699–703.

Ohlemiller, K.K., 2006. Contributions of mouse models to understanding of age-and noise-related hearing loss. Brain research 1091(1), 89–102.

Ohlemiller, K.K., Frisina, R.D., 2008. Age-related hearing loss and its cellular and molecular bases, Auditory trauma, protection, and repair. Springer, pp. 145–194.

Oliver, D.L., Huerta, M.F., 1992. Inferior and superior colliculi. The mammalian auditory pathway: neuroanatomy, 168–221.

Oliver, D.L., Morest, D.K., 1984. The central nucleus of the inferior colliculus in the cat. J Comp Neurol 222(2), 237–264.

Ouda, L., Syka, J., 2012. Immunocytochemical profiles of inferior colliculus neurons in the rat and their changes with aging. Front Neural Circuits 6, 68.

Palombi, P.S., Backoff, P.M., Caspary, D.M., 2001. Responses of young and aged rat inferior colliculus neurons to sinusoidally amplitude modulated stimuli. Hearing research 153(1-2), 174–180.

Park, C.R., Willott, J.F., Walton, J.P., 2024. Age-related changes of auditory sensitivity across the life span of CBA/CaJ mice. Hearing research 441, 108921.

Parthasarathy, A., Bartlett, E.L., 2011. Age-related auditory deficits in temporal processing in F- 344 rats. Neuroscience 192, 619–630.

Parthasarathy, A., Herrmann, B., Bartlett, E.L., 2019. Aging alters envelope representations of speech-like sounds in the inferior colliculus. Neurobiol Aging 73, 30–40.

Patel, C.R., Redhead, C., Cervi, A.L., Zhang, H., 2012. Neural sensitivity to novel sounds in the rat’s dorsal cortex of the inferior colliculus as revealed by evoked local field potentials. Hear Res 286(1-2), 41–54.

Paxinos, G., Franklin, B., 2007. The Mouse Brain in Stereotaxic Coordinates.

Peelle, J.E., Wingfield, A., 2016. The Neural Consequences of Age-Related Hearing Loss. Trends Neurosci 39(7), 486–497.

Rabang, C.F., Parthasarathy, A., Venkataraman, Y., Fisher, Z.L., Gardner, S.M., Bartlett, E.L., 2012. A computational model of inferior colliculus responses to amplitude modulated sounds in young and aged rats. Frontiers in neural circuits 6, 77.

Resnik, J., Polley, D.B., 2021. Cochlear neural degeneration disrupts hearing in background noise by increasing auditory cortex internal noise. Neuron 109(6), 984–996 e984.

Rumschlag, J.A., McClaskey, C.M., Dias, J.W., Kerouac, L.B., Noble, K.V., Panganiban, C., Lang, H., Harris, K.C., 2022. Age-related central gain with degraded neural synchrony in the auditory brainstem of mice and humans. Neurobiology of Aging 115, 50–59.

Schacht, J., Hawkins, J.E., 2005. Sketches of otohistory. Part 9: presby[a]cusis. Audiol Neurootol 10(5), 243-247.

Schofield, B.R., Cant, N.B., 1999. Descending auditory pathways: Projections from the inferior colliculus contact superior olivary cells that project bilaterally to the cochlear nuclei. Journal of Comparative Neurology 409(2), 210–223.

Schofield, B.R., Coomes, D.L., 2005. Projections from auditory cortex contact cells in the cochlear nucleus that project to the inferior colliculus. Hear Res 206(1-2), 3–11.

Schreiner, C.E., Langner, G., 1988. Periodicity coding in the inferior colliculus of the cat. II. Topographical organization. J Neurophysiol 60(6), 1823–1840.

Schreiner, C.E., Winer, J.A., 2005. The inferior colliculus. Springer.

Scott, L., Brecht, E., Philpo, A., Iyer, S., Wu, N., Mihic, S., Aldrich, R., Pierce, J., Walton, J., 2017. A novel BK channel-targeted peptide suppresses sound evoked activity in the mouse inferior colliculus. Scientific Reports 7(1), 42433.

Semple, M.N., Aitkin, L.M., Calford, M.B., Pettigrew, J.D., Phillips, D.P., 1983. Spatial receptive fields in the cat inferior colliculus. Hear Res 10(2), 203–215.

Stiebler, I., Ehret, G., 1985. Inferior colliculus of the house mouse. I. A quantitative study of tonotopic organization, frequency representation, and tone-threshold distribution. J Comp Neurol 238(1), 65–76.

Sutter, M.L., 2000. Shapes and level tolerances of frequency tuning curves in primary auditory cortex: quantitative measures and population codes. Journal of neurophysiology 84(2), 1012–1025.

Syka, J., 2020. Age-related changes in the auditory brainstem and inferior colliculus. Aging and Hearing: Causes and Consequences, 67–96.

Tallon-Baudry, C., Bertrand, O., Delpuech, C., Pernier, J., 1996. Stimulus specificity of phase-locked and non-phase-locked 40 Hz visual responses in human. Journal of Neuroscience 16(13), 4240–4249.

Tremblay, K.L., Piskosz, M., Souza, P., 2003. Effects of age and age-related hearing loss on the neural representation of speech cues. Clin Neurophysiol 114(7), 1332–1343.

Walton, J.P., Frisina, R.D., O’Neill, W.E., 1998. Age-related alteration in processing of temporal sound features in the auditory midbrain of the CBA mouse. J Neurosci 18(7), 2764–2776.

Wang, J., Ding, D., Salvi, R.J., 2002. Functional reorganization in chinchilla inferior colliculus associated with chronic and acute cochlear damage. Hear Res 168(1-2), 238–249.

Willott, J.F., Parham, K., Hunter, K.P., 1991. Comparison of the auditory sensitivity of neurons in the cochlear nucleus and inferior colliculus of young and aging C57BL/6J and CBA/J mice. Hearing research 53(1), 78–94.

Xiong, B., Alkharabsheh, A., Manohar, S., Chen, G.D., Yu, N., Zhao, X., Salvi, R., Sun, W., 2017. Hyperexcitability of inferior colliculus and acoustic startle reflex with age-related hearing loss. Hear Res 350, 32–42.

