## Supplemental Statistical Analyses for "Age-related hyperexcitability in the mouse inferior colliculus: evidence from sound-evoked local field potentials"

Dimitri L. Brunelle

#### Data Preparation

*Table S1.1: Sample Sizes by Sex and Age Group*

| Age Group | Male | Female | Animals | Channels |
|---|---|---|---|---|
| Young | 1 | 6 | 7 | 82 |
| Middle | 1 | 6 | 7 | 85 |
| Old | 15 | 5 | 20 | 243 |
| Oldest old | 5 | 2 | 7 | 85 |

*Table S1.2: Sample Sizes by Region*

| Region | Animals | Channels |
|---|---|---|
| Ventral | 41 | 158 |
| Medial | 41 | 234 |
| Dorsal | 29 | 103 |

*Table S1.3: Sample Sizes by Age Group and Region*

| Age Group | Region | Animals | Channels |
|---|---|---|---|
| Young | Ventral | 7 | 25 |
| Young | Medial | 7 | 41 |
| Young | Dorsal | 5 | 16 |
| Middle | Ventral | 7 | 26 |
| Middle | Medial | 7 | 42 |

| Age Group | Region | Animals | Channels |
|---|---|---|---|
| Middle | Dorsal | 5 | 17 |
| Old | Ventral | 20 | 82 |
| Old | Medial | 20 | 111 |
| Old | Dorsal | 15 | 50 |
| Oldest old | Ventral | 7 | 25 |
| Oldest old | Medial | 7 | 40 |
| Oldest old | Dorsal | 4 | 20 |

### Figure 1: Best Frequency and Minimum Threshold Analysis

#### Figure 1A: BF vs MT Correlation

##### Overall Correlation

*Table S2.1: Overall BF-MT Correlation*

| Test | r | 95% CI Lower | 95% CI Upper | p-value |
|---|---|---|---|---|
| Pearson Correlation | 0.184 | 0.097 | 0.268 | <0.001 |

##### Correlation by Age Group

*Table S2.2: BF-MT Correlations by Age Group*

| Age Group | n | r | 95% CI Lower | 95% CI Upper | p-value |
|---|---|---|---|---|---|
| Young | 82 | 0.206 | -0.012 | 0.405 | 0.0638 |
| Middle | 85 | 0.376 | 0.177 | 0.545 | <0.001 |
| Old | 243 | 0.333 | 0.217 | 0.441 | <0.001 |
| Oldest old | 85 | -0.098 | -0.305 | 0.117 | 0.3712 |

### Figure 1B: Best Frequency Distribution by Age and Region

#### Test for Regional Differences in BF

*Table S2.3: Kruskal-Wallis Test - BF by Region*

| Test | Factor | H | df | p-value | Sig |
|---|---|---|---|---|---|
| Kruskal-Wallis | Region | 75.125 | 2 | <0.001 | *** |

*Table S2.4: Post-hoc Pairwise Comparisons - BF by Region (Holm-Bonferroni correction)*

| Group 1 | Group 2 | W Statistic | p | p (adj) | Significance |
|---|---|---|---|---|---|
| Ventral | Medial | 24384.5000 | 0.0000 | 0.0000 | **** |
| Ventral | Dorsal | 12991.5000 | 0.0000 | 0.0000 | **** |
| Medial | Dorsal | 16232.0000 | 0.0000 | 0.0000 | **** |

#### Descriptive Statistics: BF by Region

*Table S2.5: Descriptive Statistics - BF by Region*

| Region | n | Mean (kHz) | SD | Median (kHz) | Q25 | Q75 | Min | Max |
|---|---|---|---|---|---|---|---|---|
| Ventral | 158 | 30.50 | 18.00 | 22.78 | 13.25 | 48.47 | 4.49 | 60.16 |
| Medial | 234 | 19.93 | 12.29 | 16.45 | 11.36 | 24.65 | 4.45 | 63.49 |
| Dorsal | 103 | 13.58 | 6.63 | 11.32 | 9.52 | 15.91 | 4.00 | 34.54 |

#### Test for Age Effect on BF Distribution

*Table S2.6: Kruskal-Wallis Test - BF by Age Group*

| Test | Factor | H | df | p-value | Sig |
|---|---|---|---|---|---|
| Kruskal-Wallis | Age Group | 5.321 | 3 | 0.15 | ns |

No significant age effect on BF; post-hoc tests not performed.

#### Descriptive Statistics: BF by Age and Region

*Table S2.7: Descriptive Statistics - BF by Age Group and Region*

| Age Group | Region | n | Mean (kHz) | SD | Median (kHz) | Q25 | Q75 |
|---|---|---|---|---|---|---|---|
| Young | Ventral | 25 | 31.27 | 19.50 | 36.86 | 11.77 | 48.58 |

| Age Group | Region | n | Mean (kHz) | SD | Median (kHz) | Q25 | Q75 |
|---|---|---|---|---|---|---|---|
| Young | Medial | 41 | 25.18 | 16.06 | 19.82 | 12.48 | 35.19 |
| Young | Dorsal | 16 | 14.51 | 6.68 | 12.81 | 8.83 | 19.48 |
| Middle | Ventral | 26 | 36.08 | 18.29 | 38.64 | 18.24 | 55.12 |
| Middle | Medial | 42 | 20.57 | 12.35 | 17.08 | 12.57 | 23.16 |
| Middle | Dorsal | 17 | 12.37 | 2.55 | 11.50 | 10.73 | 12.59 |
| Old | Ventral | 82 | 30.40 | 17.95 | 20.42 | 13.88 | 48.85 |
| Old | Medial | 111 | 17.97 | 10.51 | 14.61 | 11.02 | 19.82 |
| Old | Dorsal | 50 | 12.00 | 5.67 | 11.12 | 8.20 | 13.08 |
| Oldest old | Ventral | 25 | 24.24 | 15.10 | 15.56 | 11.46 | 37.91 |
| Oldest old | Medial | 40 | 19.29 | 11.18 | 13.36 | 10.91 | 25.14 |
| Oldest old | Dorsal | 20 | 17.79 | 9.22 | 10.83 | 10.22 | 23.14 |

#### Two-Way Test: Age × Region Effect on BF

*Table S2.8: Two-Way ANOVA - Age × Region Effect on BF*

| Factor | df | Sum Sq | Mean Sq | F | p-value | Sig |
|---|---|---|---|---|---|---|
| age.cat | 3.000 | 1449.314 | 483.105 | 2.694 | 0.0455 | * |
| region | 2.000 | 19804.166 | 9902.083 | 55.216 | <0.001 | *** |
| age.cat:region | 6.000 | 2387.444 | 397.907 | 2.219 | 0.0402 | * |
| Residuals | 483.000 | 86618.059 | 179.333 | | NA | ns |

*Table S2.9: Effect Sizes - Age × Region on BF*

| Factor | $\eta^2$ | Partial $\eta^2$ |
|---|---|---|
| age.cat | 0.013 | 0.016 |
| region | 0.180 | 0.186 |
| age.cat:region | 0.022 | 0.027 |

*Table S2.10: Levene's Test for Homogeneity of Variance*

| Test | F | df1 | df2 | p-value |
|---|---|---|---|---|

| Test | F | df1 | df2 | p-value |
|---|---|---|---|---|
| Levene's Test | 12.026 | 11 | 483 | <0.001 |

#### Test for Age Effect on MT Distribution

*Table S2.11: Kruskal-Wallis Test - MT by Age Group*

| Test | Factor | H | df | p-value | Sig |
|---|---|---|---|---|---|
| Kruskal-Wallis | Age Group | 134.37 | 3 | < 2.2e-16 | **** |

*Table S2.12: Post-hoc Pairwise Comparisons - MT by Age Group (Holm-Bonferroni correction)*

| Group 1 | Group 2 | W | p | p (adj) | Sig |
|---|---|---|---|---|---|
| Young | Middle | 3814 | 2.93e-1 | 2.93e-1 | ns |
| Young | Old | 6896 | 3.07e-5 | 6.14e-5 | **** |
| Young | Oldest old | 1020 | 2.98e-15 | 1.19e-14 | **** |
| Middle | Old | 5920 | 4.74e-9 | 1.42e-8 | **** |
| Middle | Oldest old | 645 | 2.29e-20 | 1.37e-19 | **** |
| Old | Oldest old | 3677 | 9.75e-19 | 4.88e-18 | **** |

#### Descriptive Statistics: MT by Age Group

*Table S2.13: Descriptive Statistics - MT by Age Group*

| Age Group | n | Mean (dB SPL) | SD | Median | Q25 | Q75 |
|---|---|---|---|---|---|---|
| Young | 82 | 28.9 | 18.3 | 24.2 | 16.1 | 37 |
| Middle | 85 | 25.4 | 16 | 21.9 | 16 | 31.9 |
| Old | 243 | 36.5 | 14.8 | 35 | 24.7 | 47 |
| Oldest old | 85 | 54 | 10.6 | 57 | 47 | 61.9 |

### Figure 2: LFP Amplitude Analysis

#### Normality Tests

*Table S3.1: Normality Tests (Shapiro-Wilk) by Age and Region*

| Age Group | Region | n | Shapiro p-value | Normal? |
|---|---|---|---|---|
| Young | Ventral | 25 | 0.0026 | No |
| Young | Medial | 41 | 0.1043 | Yes |
| Young | Dorsal | 16 | 0.0038 | No |
| Middle | Ventral | 26 | 0.5069 | Yes |
| Middle | Medial | 42 | 0.6968 | Yes |
| Middle | Dorsal | 17 | 0.4861 | Yes |
| Old | Ventral | 82 | 0.0154 | No |
| Old | Medial | 111 | 0.1194 | Yes |
| Old | Dorsal | 50 | 0.1989 | Yes |
| Oldest old | Ventral | 25 | 0.0498 | No |
| Oldest old | Medial | 40 | 0.1019 | Yes |
| Oldest old | Dorsal | 20 | 0.5475 | Yes |

#### Main Effect of Age by Region

*Table S3.2: Kruskal-Wallis Tests - Age Effect on N1 Amplitude by Region (80 dB SPL)*

| Region | H | df | p-value | Sig |
|---|---|---|---|---|
| Ventral | 4.035 | 3 | 0.25770 | ns |
| Medial | 14.251 | 3 | 0.00258 | ** |
| Dorsal | 26.122 | 3 | $< 0.001$ | *** |

#### Post-hoc Pairwise Comparisons by Region

*Table S3.3: Post-hoc Pairwise Comparisons (Wilcoxon, Holm-Bonferroni correction) by Region*

| Region | Group 1 | Group 2 | W | p | p (adj) | Sig |
|---|---|---|---|---|---|---|
| Ventral | Young | Middle | 399.0000 | 0.1670 | 0.8350 | ns |
| Ventral | Young | Old | 1307.0000 | 0.0380 | 0.2290 | ns |
| Ventral | Young | Oldest old | 330.0000 | 0.7440 | 1.0000 | ns |
| Ventral | Middle | Old | 1148.0000 | 0.5580 | 1.0000 | ns |
| Ventral | Middle | Oldest old | 300.0000 | 0.6470 | 1.0000 | ns |
| Ventral | Old | Oldest old | 937.0000 | 0.5190 | 1.0000 | ns |
| Medial | Young | Middle | 838.0000 | 0.8380 | 0.8380 | ns |
| Medial | Young | Old | 1948.0000 | 0.1750 | 0.3960 | ns |
| Medial | Young | Oldest old | 534.0000 | 0.0070 | 0.0290 | * |
| Medial | Middle | Old | 1962.0000 | 0.1320 | 0.3960 | ns |
| Medial | Middle | Oldest old | 467.0000 | 0.0004 | 0.0030 | ** |
| Medial | Old | Oldest old | 1566.0000 | 0.0060 | 0.0290 | * |
| Dorsal | Young | Middle | 109.0000 | 0.3450 | 0.3450 | ns |
| Dorsal | Young | Old | 215.0000 | 0.0060 | 0.0170 | * |
| Dorsal | Young | Oldest old | 41.0000 | 0.0001 | 0.0003 | *** |
| Dorsal | Middle | Old | 268.0000 | 0.0240 | 0.0480 | * |
| Dorsal | Middle | Oldest old | 37.0000 | 0.0000 | 0.0001 | **** |
| Dorsal | Old | Oldest old | 258.0000 | 0.0020 | 0.0070 | ** |

#### Descriptive Statistics

*Table S3.4: Descriptive Statistics - N1 Amplitude ($\mu V$) by Age and Region*

| Age Group | Region | n | Mean | SD | Median | Q25 | Q75 |
|---|---|---|---|---|---|---|---|
| Young | Ventral | 25 | 336.28 | 145.71 | 323.23 | 248.39 | 373.60 |

| Age Group | Region | n | Mean | SD | Median | Q25 | Q75 |
|---|---|---|---|---|---|---|---|
| Young | Medial | 41 | 300.39 | 144.98 | 296.82 | 186.08 | 379.97 |
| Young | Dorsal | 16 | 163.93 | 133.12 | 146.04 | 78.69 | 180.70 |
| Middle | Ventral | 26 | 274.46 | 104.85 | 272.14 | 184.85 | 355.55 |
| Middle | Medial | 42 | 294.04 | 103.48 | 285.01 | 229.41 | 379.17 |
| Middle | Dorsal | 17 | 170.02 | 91.08 | 177.66 | 102.47 | 242.59 |
| Old | Ventral | 82 | 268.91 | 138.24 | 256.76 | 158.36 | 358.87 |
| Old | Medial | 111 | 327.51 | 130.11 | 338.42 | 234.60 | 427.06 |
| Old | Dorsal | 50 | 264.05 | 145.92 | 239.57 | 159.94 | 360.70 |
| Oldest old | Ventral | 25 | 296.91 | 175.77 | 353.44 | 122.17 | 406.67 |
| Oldest old | Medial | 40 | 399.13 | 156.23 | 419.00 | 301.07 | 509.66 |
| Oldest old | Dorsal | 20 | 397.46 | 154.10 | 401.15 | 279.54 | 505.11 |

### Figure 3: Amplitude-Intensity Functions

#### Two-way Repeated Measures ANOVA: Age × Level by Region

*Table S4.1: Two-Way Repeated Measures ANOVA - Age × Level for N1 Amplitude*

| Region | Stratum | Factor | df1 | df2 | F | p-value | Sig | Summary |
|---|---|---|---|---|---|---|---|---|
| Ventral | Between-subjects | Age | 3.000 | 154.000 | 8.576 | <0.001 | *** | F(3, 154) = 8.576 |
| Ventral | Within-subjects | Level | 8.000 | 1232.000 | 337.028 | <0.001 | *** | F(8, 1232) = 337.028 |
| Ventral | Within-subjects | Age × Level | 24.000 | 1232.000 | 8.326 | <0.001 | *** | F(24, 1232) = 8.326 |
| Medial | Between-subjects | Age | 3.000 | 230.000 | 6.051 | <0.001 | *** | F(3, 230) = 6.051 |
| Medial | Within-subjects | Level | 8.000 | 1840.000 | 627.033 | <0.001 | *** | F(8, 1840) = 627.033 |

| Region | Stratum | Factor | df1 | df2 | F | p-value | Sig | Summary |
|---|---|---|---|---|---|---|---|---|
| Medial | Within-subjects | Age × Level | 24.000 | 1840.000 | 25.134 | <0.001 | *** | F(24, 1840) = 25.134 |
| Dorsal | Between-subjects | Age | 3.000 | 99.000 | 1.314 | 0.274 | ns | F(3, 99) = 1.314 |
| Dorsal | Within-subjects | Level | 8.000 | 792.000 | 188.196 | <0.001 | *** | F(8, 792) = 188.196 |
| Dorsal | Within-subjects | Age × Level | 24.000 | 792.000 | 12.404 | <0.001 | *** | F(24, 792) = 12.404 |

#### Descriptive Statistics by Level and Region

*Table S4.2: Descriptive Statistics - N1 Amplitude (μV) by Level, Region, and Age*

| Region | Level (dB SPL) | Age Group | n | Mean | SD | SEM |
|---|---|---|---|---|---|---|
| Ventral | 0 | Young | 25 | 8.53 | 8.42 | 1.68 |
| Ventral | 0 | Middle | 26 | 10.00 | 3.16 | 0.62 |
| Ventral | 0 | Old | 82 | 8.79 | 8.03 | 0.89 |
| Ventral | 0 | Oldest old | 25 | 14.01 | 10.26 | 2.05 |
| Ventral | 10 | Young | 25 | 13.01 | 7.84 | 1.57 |
| Ventral | 10 | Middle | 26 | 10.71 | 12.96 | 2.54 |
| Ventral | 10 | Old | 82 | 7.77 | 6.46 | 0.71 |
| Ventral | 10 | Oldest old | 25 | 8.63 | 8.23 | 1.65 |
| Ventral | 20 | Young | 25 | 34.65 | 29.22 | 5.84 |
| Ventral | 20 | Middle | 26 | 27.91 | 33.20 | 6.51 |
| Ventral | 20 | Old | 82 | 10.97 | 15.35 | 1.69 |
| Ventral | 20 | Oldest old | 25 | 9.89 | 7.77 | 1.55 |
| Ventral | 30 | Young | 25 | 186.25 | 61.50 | 12.30 |
| Ventral | 30 | Middle | 26 | 123.82 | 121.53 | 23.83 |
| Ventral | 30 | Old | 82 | 32.01 | 45.59 | 5.03 |

| Region | Level (dB SPL) | Age Group | n | Mean | SD | SEM |
|---|---|---|---|---|---|---|
| Ventral | 30 | Oldest old | 25 | 14.27 | 10.81 | 2.16 |
| Ventral | 40 | Young | 25 | 260.92 | 111.94 | 22.39 |
| Ventral | 40 | Middle | 26 | 235.55 | 156.96 | 30.78 |
| Ventral | 40 | Old | 82 | 95.64 | 74.83 | 8.26 |
| Ventral | 40 | Oldest old | 25 | 18.67 | 16.13 | 3.23 |
| Ventral | 50 | Young | 25 | 287.15 | 130.99 | 26.20 |
| Ventral | 50 | Middle | 26 | 261.30 | 148.88 | 29.20 |
| Ventral | 50 | Old | 82 | 195.66 | 136.20 | 15.04 |
| Ventral | 50 | Oldest old | 25 | 114.66 | 155.39 | 31.08 |
| Ventral | 60 | Young | 25 | 301.69 | 139.28 | 27.86 |
| Ventral | 60 | Middle | 26 | 257.44 | 126.66 | 24.84 |
| Ventral | 60 | Old | 82 | 248.62 | 144.04 | 15.91 |
| Ventral | 60 | Oldest old | 25 | 214.49 | 183.31 | 36.66 |
| Ventral | 70 | Young | 25 | 313.88 | 134.02 | 26.80 |
| Ventral | 70 | Middle | 26 | 271.77 | 113.82 | 22.32 |
| Ventral | 70 | Old | 82 | 262.67 | 146.38 | 16.17 |
| Ventral | 70 | Oldest old | 25 | 284.13 | 189.39 | 37.88 |
| Ventral | 80 | Young | 25 | 336.28 | 145.71 | 29.14 |
| Ventral | 80 | Middle | 26 | 274.46 | 104.85 | 20.56 |
| Ventral | 80 | Old | 82 | 268.91 | 138.24 | 15.27 |
| Ventral | 80 | Oldest old | 25 | 296.91 | 175.77 | 35.15 |
| Medial | 0 | Young | 41 | 6.64 | 5.31 | 0.83 |
| Medial | 0 | Middle | 42 | 13.02 | 5.59 | 0.86 |
| Medial | 0 | Old | 111 | 8.65 | 6.80 | 0.65 |
| Medial | 0 | Oldest old | 40 | 9.72 | 7.80 | 1.23 |

| Region | Level (dB SPL) | Age Group | n | Mean | SD | SEM |
|---|---|---|---|---|---|---|
| Medial | 10 | Young | 41 | 17.93 | 12.05 | 1.88 |
| Medial | 10 | Middle | 42 | 14.86 | 12.76 | 1.97 |
| Medial | 10 | Old | 111 | 8.52 | 5.91 | 0.56 |
| Medial | 10 | Oldest old | 40 | 7.13 | 6.37 | 1.01 |
| Medial | 20 | Young | 41 | 72.10 | 64.08 | 10.01 |
| Medial | 20 | Middle | 42 | 62.89 | 79.98 | 12.34 |
| Medial | 20 | Old | 111 | 15.03 | 20.01 | 1.90 |
| Medial | 20 | Oldest old | 40 | 8.28 | 5.94 | 0.94 |
| Medial | 30 | Young | 41 | 231.61 | 94.13 | 14.70 |
| Medial | 30 | Middle | 42 | 201.51 | 124.80 | 19.26 |
| Medial | 30 | Old | 111 | 69.75 | 95.96 | 9.11 |
| Medial | 30 | Oldest old | 40 | 9.51 | 4.78 | 0.76 |
| Medial | 40 | Young | 41 | 295.28 | 117.09 | 18.29 |
| Medial | 40 | Middle | 42 | 314.48 | 114.76 | 17.71 |
| Medial | 40 | Old | 111 | 184.40 | 163.89 | 15.56 |
| Medial | 40 | Oldest old | 40 | 22.33 | 17.55 | 2.77 |
| Medial | 50 | Young | 41 | 285.41 | 124.28 | 19.41 |
| Medial | 50 | Middle | 42 | 310.73 | 108.18 | 16.69 |
| Medial | 50 | Old | 111 | 285.31 | 161.39 | 15.32 |
| Medial | 50 | Oldest old | 40 | 138.97 | 166.31 | 26.30 |
| Medial | 60 | Young | 41 | 293.82 | 132.38 | 20.67 |
| Medial | 60 | Middle | 42 | 299.55 | 107.58 | 16.60 |
| Medial | 60 | Old | 111 | 333.92 | 150.14 | 14.25 |
| Medial | 60 | Oldest old | 40 | 288.28 | 167.05 | 26.41 |
| Medial | 70 | Young | 41 | 295.18 | 138.27 | 21.59 |

| Region | Level (dB SPL) | Age Group | n | Mean | SD | SEM |
|---|---|---|---|---|---|---|
| Medial | 70 | Middle | 42 | 299.00 | 104.15 | 16.07 |
| Medial | 70 | Old | 111 | 331.84 | 143.52 | 13.62 |
| Medial | 70 | Oldest old | 40 | 370.47 | 162.16 | 25.64 |
| Medial | 80 | Young | 41 | 300.39 | 144.98 | 22.64 |
| Medial | 80 | Middle | 42 | 294.04 | 103.48 | 15.97 |
| Medial | 80 | Old | 111 | 327.51 | 130.11 | 12.35 |
| Medial | 80 | Oldest old | 40 | 399.13 | 156.23 | 24.70 |
| Dorsal | 0 | Young | 16 | 5.34 | 4.93 | 1.23 |
| Dorsal | 0 | Middle | 17 | 9.97 | 4.12 | 1.00 |
| Dorsal | 0 | Old | 50 | 7.82 | 4.51 | 0.64 |
| Dorsal | 0 | Oldest old | 20 | 7.47 | 3.78 | 0.85 |
| Dorsal | 10 | Young | 16 | 15.29 | 9.74 | 2.43 |
| Dorsal | 10 | Middle | 17 | 10.34 | 6.41 | 1.55 |
| Dorsal | 10 | Old | 50 | 7.40 | 5.24 | 0.74 |
| Dorsal | 10 | Oldest old | 20 | 6.70 | 4.44 | 0.99 |
| Dorsal | 20 | Young | 16 | 59.18 | 39.34 | 9.84 |
| Dorsal | 20 | Middle | 17 | 45.69 | 57.14 | 13.86 |
| Dorsal | 20 | Old | 50 | 13.10 | 13.94 | 1.97 |
| Dorsal | 20 | Oldest old | 20 | 7.68 | 6.01 | 1.34 |
| Dorsal | 30 | Young | 16 | 131.34 | 48.20 | 12.05 |
| Dorsal | 30 | Middle | 17 | 115.04 | 83.54 | 20.26 |
| Dorsal | 30 | Old | 50 | 74.41 | 71.34 | 10.09 |
| Dorsal | 30 | Oldest old | 20 | 8.45 | 5.95 | 1.33 |
| Dorsal | 40 | Young | 16 | 147.56 | 60.78 | 15.19 |
| Dorsal | 40 | Middle | 17 | 165.50 | 82.50 | 20.01 |

| Region | Level (dB SPL) | Age Group | n | Mean | SD | SEM |
|---|---|---|---|---|---|---|
| Dorsal | 40 | Old | 50 | 166.60 | 121.05 | 17.12 |
| Dorsal | 40 | Oldest old | 20 | 23.46 | 18.18 | 4.06 |
| Dorsal | 50 | Young | 16 | 146.35 | 72.18 | 18.05 |
| Dorsal | 50 | Middle | 17 | 171.47 | 87.76 | 21.29 |
| Dorsal | 50 | Old | 50 | 224.51 | 142.67 | 20.18 |
| Dorsal | 50 | Oldest old | 20 | 178.94 | 180.88 | 40.45 |
| Dorsal | 60 | Young | 16 | 160.90 | 98.04 | 24.51 |
| Dorsal | 60 | Middle | 17 | 167.66 | 91.02 | 22.08 |
| Dorsal | 60 | Old | 50 | 254.25 | 158.16 | 22.37 |
| Dorsal | 60 | Oldest old | 20 | 315.88 | 192.46 | 43.04 |
| Dorsal | 70 | Young | 16 | 169.83 | 120.78 | 30.20 |
| Dorsal | 70 | Middle | 17 | 171.98 | 90.41 | 21.93 |
| Dorsal | 70 | Old | 50 | 260.66 | 153.11 | 21.65 |
| Dorsal | 70 | Oldest old | 20 | 350.71 | 168.29 | 37.63 |
| Dorsal | 80 | Young | 16 | 163.93 | 133.12 | 33.28 |
| Dorsal | 80 | Middle | 17 | 170.02 | 91.08 | 22.09 |
| Dorsal | 80 | Old | 50 | 264.05 | 145.92 | 20.64 |
| Dorsal | 80 | Oldest old | 20 | 397.46 | 154.10 | 34.46 |

#### Post-hoc Tests: Age Effect at Each Level by Region

*Table S4.3: Kruskal-Wallis Tests - Age Effect at Each Level by Region*

| Region | Level (dB SPL) | H | df | p-value | Sig |
|---|---|---|---|---|---|
| Ventral | 0 | 12.425 | 3 | 0.00606 | ** |
| Ventral | 10 | 9.435 | 3 | 0.024 | * |
| Ventral | 20 | 25.407 | 3 | <0.001 | *** |

| Region | Level (dB SPL) | H | df | p-value | Sig |
|---|---|---|---|---|---|
| Ventral | 30 | 85.847 | 3 | <0.001 | *** |
| Ventral | 40 | 87.535 | 3 | <0.001 | *** |
| Ventral | 50 | 28.460 | 3 | <0.001 | *** |
| Ventral | 60 | 6.953 | 3 | 0.0734 | ns |
| Ventral | 70 | 3.001 | 3 | 0.391 | ns |
| Ventral | 80 | 4.035 | 3 | 0.258 | ns |
| Medial | 0 | 25.585 | 3 | <0.001 | *** |
| Medial | 10 | 28.685 | 3 | <0.001 | *** |
| Medial | 20 | 70.508 | 3 | <0.001 | *** |
| Medial | 30 | 124.710 | 3 | <0.001 | *** |
| Medial | 40 | 111.762 | 3 | <0.001 | *** |
| Medial | 50 | 40.089 | 3 | <0.001 | *** |
| Medial | 60 | 5.086 | 3 | 0.166 | ns |
| Medial | 70 | 7.436 | 3 | 0.0592 | ns |
| Medial | 80 | 14.251 | 3 | 0.00258 | ** |
| Dorsal | 0 | 9.840 | 3 | 0.02 | * |
| Dorsal | 10 | 9.373 | 3 | 0.0247 | * |
| Dorsal | 20 | 20.185 | 3 | <0.001 | *** |
| Dorsal | 30 | 43.857 | 3 | <0.001 | *** |
| Dorsal | 40 | 34.175 | 3 | <0.001 | *** |
| Dorsal | 50 | 5.509 | 3 | 0.138 | ns |
| Dorsal | 60 | 10.671 | 3 | 0.0136 | * |
| Dorsal | 70 | 17.242 | 3 | <0.001 | *** |
| Dorsal | 80 | 26.122 | 3 | <0.001 | *** |

*Table S4.4: Pairwise Wilcoxon Tests - N1 Amplitude at Each Level by Region (Holm-Bonferroni correction)*

| Region | Level (dB SPL) | Group 1 | Group 2 | W | p | p (adj) | Sig |
|---|---|---|---|---|---|---|---|
| Ventral | 0 | Middle | Old | 1426.000 | 0.010000 | 0.059000 | ns |
| Ventral | 0 | Middle | Oldest old | 276.000 | 0.364000 | 0.728000 | ns |
| Ventral | 0 | Old | Oldest old | 680.000 | 0.011000 | 0.059000 | ns |
| Ventral | 0 | Young | Middle | 202.000 | 0.020000 | 0.080000 | ns |
| Ventral | 0 | Young | Old | 994.000 | 0.822000 | 0.822000 | ns |
| Ventral | 0 | Young | Oldest old | 198.000 | 0.026000 | 0.080000 | ns |
| Ventral | 10 | Middle | Old | 1026.000 | 0.777000 | 1.000000 | ns |
| Ventral | 10 | Middle | Oldest old | 311.000 | 0.801000 | 1.000000 | ns |
| Ventral | 10 | Old | Oldest old | 1010.000 | 0.915000 | 1.000000 | ns |
| Ventral | 10 | Young | Middle | 433.000 | 0.042000 | 0.168000 | ns |
| Ventral | 10 | Young | Old | 1435.000 | 0.003000 | 0.015000 | * |
| Ventral | 10 | Young | Oldest old | 438.000 | 0.014000 | 0.072000 | ns |
| Ventral | 20 | Middle | Old | 1029.000 | 0.793000 | 1.000000 | ns |
| Ventral | 20 | Middle | Oldest old | 333.000 | 0.889000 | 1.000000 | ns |
| Ventral | 20 | Old | Oldest old | 970.000 | 0.688000 | 1.000000 | ns |
| Ventral | 20 | Young | Middle | 428.000 | 0.053000 | 0.212000 | ns |
| Ventral | 20 | Young | Old | 1755.000 | $< 1e-04$ | $< 1e-04$ | **** |
| Ventral | 20 | Young | Oldest old | 534.000 | $< 1e-04$ | $< 1e-04$ | **** |
| Ventral | 30 | Middle | Old | 1883.000 | $< 1e-04$ | $< 1e-04$ | **** |
| Ventral | 30 | Middle | Oldest old | 636.000 | $< 1e-04$ | $< 1e-04$ | **** |
| Ventral | 30 | Old | Oldest old | 1335.000 | 0.023000 | 0.023000 | * |
| Ventral | 30 | Young | Middle | 498.000 | 0.000858 | 0.002000 | ** |

*Table S4.4: Pairwise Wilcoxon Tests - N1 Amplitude at Each Level by Region (Holm-Bonferroni correction)*

| Region | Level (dB SPL) | Group 1 | Group 2 | W | p | p (adj) | Sig |
|---|---|---|---|---|---|---|---|
| Ventral | 30 | Young | Old | 1986.000 | < 1e-04 | < 1e-04 | **** |
| Ventral | 30 | Young | Oldest old | 624.000 | < 1e-04 | < 1e-04 | **** |
| Ventral | 40 | Middle | Old | 1769.000 | < 1e-04 | < 1e-04 | **** |
| Ventral | 40 | Middle | Oldest old | 649.000 | < 1e-04 | < 1e-04 | **** |
| Ventral | 40 | Old | Oldest old | 1841.000 | < 1e-04 | < 1e-04 | **** |
| Ventral | 40 | Young | Middle | 406.000 | 0.130000 | 0.130000 | ns |
| Ventral | 40 | Young | Old | 1842.000 | < 1e-04 | < 1e-04 | **** |
| Ventral | 40 | Young | Oldest old | 624.000 | < 1e-04 | < 1e-04 | **** |
| Ventral | 50 | Middle | Old | 1371.000 | 0.029000 | 0.057000 | ns |
| Ventral | 50 | Middle | Oldest old | 521.000 | 0.000133 | 0.000665 | *** |
| Ventral | 50 | Old | Oldest old | 1515.000 | 0.000314 | 0.001000 | ** |
| Ventral | 50 | Young | Middle | 385.000 | 0.265000 | 0.265000 | ns |
| Ventral | 50 | Young | Old | 1475.000 | 0.000935 | 0.003000 | ** |
| Ventral | 50 | Young | Oldest old | 510.000 | < 1e-04 | 0.000404 | *** |
| Ventral | 60 | Middle | Old | 1138.000 | 0.607000 | 0.607000 | ns |
| Ventral | 60 | Middle | Oldest old | 404.000 | 0.140000 | 0.476000 | ns |
| Ventral | 60 | Old | Oldest old | 1237.000 | 0.119000 | 0.476000 | ns |
| Ventral | 60 | Young | Middle | 388.000 | 0.241000 | 0.482000 | ns |
| Ventral | 60 | Young | Old | 1281.000 | 0.060000 | 0.300000 | ns |
| Ventral | 60 | Young | Oldest old | 427.000 | 0.026000 | 0.157000 | ns |
| Ventral | 70 | Middle | Old | 1165.000 | 0.479000 | 1.000000 | ns |
| Ventral | 70 | Middle | Oldest old | 323.000 | 0.978000 | 1.000000 | ns |
| Ventral | 70 | Old | Oldest old | 986.000 | 0.777000 | 1.000000 | ns |

*Table S4.4: Pairwise Wilcoxon Tests - N1 Amplitude at Each Level by Region (Holm-Bonferroni correction)*

| Region | Level (dB SPL) | Group 1 | Group 2 | W | p | p (adj) | Sig |
|---|---|---|---|---|---|---|---|
| Ventral | 70 | Young | Middle | 380.000 | 0.307000 | 1.000000 | ns |
| Ventral | 70 | Young | Old | 1275.000 | 0.066000 | 0.397000 | ns |
| Ventral | 70 | Young | Oldest old | 338.000 | 0.631000 | 1.000000 | ns |
| Ventral | 80 | Middle | Old | 1148.000 | 0.558000 | 1.000000 | ns |
| Ventral | 80 | Middle | Oldest old | 300.000 | 0.647000 | 1.000000 | ns |
| Ventral | 80 | Old | Oldest old | 937.000 | 0.519000 | 1.000000 | ns |
| Ventral | 80 | Young | Middle | 399.000 | 0.167000 | 0.835000 | ns |
| Ventral | 80 | Young | Old | 1307.000 | 0.038000 | 0.229000 | ns |
| Ventral | 80 | Young | Oldest old | 330.000 | 0.744000 | 1.000000 | ns |
| Medial | 0 | Middle | Old | 3354.000 | < 1e-04 | 0.000146 | *** |
| Medial | 0 | Middle | Oldest old | 1143.000 | 0.005000 | 0.018000 | * |
| Medial | 0 | Old | Oldest old | 2092.000 | 0.591000 | 0.591000 | ns |
| Medial | 0 | Young | Middle | 323.000 | < 1e-04 | < 1e-04 | **** |
| Medial | 0 | Young | Old | 1903.000 | 0.123000 | 0.265000 | ns |
| Medial | 0 | Young | Oldest old | 639.000 | 0.088000 | 0.265000 | ns |
| Medial | 10 | Middle | Old | 2829.000 | 0.042000 | 0.126000 | ns |
| Medial | 10 | Middle | Oldest old | 1119.000 | 0.009000 | 0.037000 | * |
| Medial | 10 | Old | Oldest old | 2643.000 | 0.075000 | 0.150000 | ns |
| Medial | 10 | Young | Middle | 1023.000 | 0.142000 | 0.150000 | ns |
| Medial | 10 | Young | Old | 3376.000 | < 1e-04 | < 1e-04 | **** |
| Medial | 10 | Young | Oldest old | 1307.000 | < 1e-04 | < 1e-04 | **** |
| Medial | 20 | Middle | Old | 3297.000 | < 1e-04 | 0.000237 | *** |
| Medial | 20 | Middle | Oldest old | 1267.000 | < 1e-04 | 0.000196 | *** |

*Table S4.4: Pairwise Wilcoxon Tests - N1 Amplitude at Each Level by Region (Holm-Bonferroni correction)*

| Region | Level (dB SPL) | Group 1 | Group 2 | W | p | p (adj) | Sig |
|---|---|---|---|---|---|---|---|
| Medial | 20 | Old | Oldest old | 2640.000 | 0.077000 | 0.077000 | ns |
| Medial | 20 | Young | Middle | 1131.000 | 0.014000 | 0.027000 | * |
| Medial | 20 | Young | Old | 4022.000 | < 1e-04 | < 1e-04 | **** |
| Medial | 20 | Young | Oldest old | 1548.000 | < 1e-04 | < 1e-04 | **** |
| Medial | 30 | Middle | Old | 3924.000 | < 1e-04 | < 1e-04 | **** |
| Medial | 30 | Middle | Oldest old | 1680.000 | < 1e-04 | < 1e-04 | **** |
| Medial | 30 | Old | Oldest old | 3543.000 | < 1e-04 | < 1e-04 | **** |
| Medial | 30 | Young | Middle | 1016.000 | 0.160000 | 0.160000 | ns |
| Medial | 30 | Young | Old | 4070.000 | < 1e-04 | < 1e-04 | **** |
| Medial | 30 | Young | Oldest old | 1640.000 | < 1e-04 | < 1e-04 | **** |
| Medial | 40 | Middle | Old | 3619.000 | < 1e-04 | < 1e-04 | **** |
| Medial | 40 | Middle | Oldest old | 1680.000 | < 1e-04 | < 1e-04 | **** |
| Medial | 40 | Old | Oldest old | 4057.000 | < 1e-04 | < 1e-04 | **** |
| Medial | 40 | Young | Middle | 772.000 | 0.422000 | 0.422000 | ns |
| Medial | 40 | Young | Old | 3424.000 | < 1e-04 | < 1e-04 | **** |
| Medial | 40 | Young | Oldest old | 1640.000 | < 1e-04 | < 1e-04 | **** |
| Medial | 50 | Middle | Old | 2687.000 | 0.146000 | 0.438000 | ns |
| Medial | 50 | Middle | Oldest old | 1425.000 | < 1e-04 | < 1e-04 | **** |
| Medial | 50 | Old | Oldest old | 3518.000 | < 1e-04 | < 1e-04 | **** |
| Medial | 50 | Young | Middle | 753.000 | 0.329000 | 0.658000 | ns |
| Medial | 50 | Young | Old | 2352.000 | 0.752000 | 0.752000 | ns |
| Medial | 50 | Young | Oldest old | 1344.000 | < 1e-04 | < 1e-04 | **** |
| Medial | 60 | Middle | Old | 2018.000 | 0.201000 | 0.804000 | ns |

*Table S4.4: Pairwise Wilcoxon Tests - N1 Amplitude at Each Level by Region (Holm-Bonferroni correction)*

| Region | Level (dB SPL) | Group 1 | Group 2 | W | p | p (adj) | Sig |
|---|---|---|---|---|---|---|---|
| Medial | 60 | Middle | Oldest old | 937.000 | 0.373000 | 1.000000 | ns |
| Medial | 60 | Old | Oldest old | 2661.000 | 0.063000 | 0.379000 | ns |
| Medial | 60 | Young | Middle | 833.000 | 0.803000 | 1.000000 | ns |
| Medial | 60 | Young | Old | 1925.000 | 0.146000 | 0.730000 | ns |
| Medial | 60 | Young | Oldest old | 884.000 | 0.551000 | 1.000000 | ns |
| Medial | 70 | Middle | Old | 1984.000 | 0.157000 | 0.528000 | ns |
| Medial | 70 | Middle | Oldest old | 597.000 | 0.024000 | 0.143000 | ns |
| Medial | 70 | Old | Oldest old | 1909.000 | 0.190000 | 0.528000 | ns |
| Medial | 70 | Young | Middle | 821.000 | 0.720000 | 0.720000 | ns |
| Medial | 70 | Young | Old | 1912.000 | 0.132000 | 0.528000 | ns |
| Medial | 70 | Young | Oldest old | 586.000 | 0.027000 | 0.143000 | ns |
| Medial | 80 | Middle | Old | 1962.000 | 0.132000 | 0.396000 | ns |
| Medial | 80 | Middle | Oldest old | 467.000 | 0.000435 | 0.003000 | ** |
| Medial | 80 | Old | Oldest old | 1566.000 | 0.006000 | 0.029000 | * |
| Medial | 80 | Young | Middle | 838.000 | 0.838000 | 0.838000 | ns |
| Medial | 80 | Young | Old | 1948.000 | 0.175000 | 0.396000 | ns |
| Medial | 80 | Young | Oldest old | 534.000 | 0.007000 | 0.029000 | * |
| Dorsal | 0 | Middle | Old | 564.000 | 0.046000 | 0.206000 | ns |
| Dorsal | 0 | Middle | Oldest old | 231.000 | 0.065000 | 0.206000 | ns |
| Dorsal | 0 | Old | Oldest old | 519.000 | 0.810000 | 0.810000 | ns |
| Dorsal | 0 | Young | Middle | 62.000 | 0.007000 | 0.041000 | * |
| Dorsal | 0 | Young | Old | 263.000 | 0.041000 | 0.206000 | ns |
| Dorsal | 0 | Young | Oldest old | 108.000 | 0.102000 | 0.206000 | ns |

*Table S4.4: Pairwise Wilcoxon Tests - N1 Amplitude at Each Level by Region (Holm-Bonferroni correction)*

| Region | Level (dB SPL) | Group 1 | Group 2 | W | p | p (adj) | Sig |
|---|---|---|---|---|---|---|---|
| Dorsal | 10 | Middle | Old | 536.000 | 0.111000 | 0.330000 | ns |
| Dorsal | 10 | Middle | Oldest old | 231.000 | 0.065000 | 0.259000 | ns |
| Dorsal | 10 | Old | Oldest old | 501.000 | 0.995000 | 0.995000 | ns |
| Dorsal | 10 | Young | Middle | 181.000 | 0.110000 | 0.330000 | ns |
| Dorsal | 10 | Young | Old | 570.000 | 0.011000 | 0.067000 | ns |
| Dorsal | 10 | Young | Oldest old | 228.000 | 0.030000 | 0.152000 | ns |
| Dorsal | 20 | Middle | Old | 538.000 | 0.105000 | 0.315000 | ns |
| Dorsal | 20 | Middle | Oldest old | 231.000 | 0.065000 | 0.259000 | ns |
| Dorsal | 20 | Old | Oldest old | 579.000 | 0.307000 | 0.552000 | ns |
| Dorsal | 20 | Young | Middle | 167.000 | 0.276000 | 0.552000 | ns |
| Dorsal | 20 | Young | Old | 679.000 | < 1e-04 | 0.000185 | *** |
| Dorsal | 20 | Young | Oldest old | 279.000 | < 1e-04 | 0.000297 | *** |
| Dorsal | 30 | Middle | Old | 552.000 | 0.068000 | 0.137000 | ns |
| Dorsal | 30 | Middle | Oldest old | 321.000 | < 1e-04 | < 1e-04 | **** |
| Dorsal | 30 | Old | Oldest old | 901.000 | < 1e-04 | < 1e-04 | **** |
| Dorsal | 30 | Young | Middle | 152.000 | 0.581000 | 0.581000 | ns |
| Dorsal | 30 | Young | Old | 618.000 | 0.001000 | 0.003000 | ** |
| Dorsal | 30 | Young | Oldest old | 320.000 | < 1e-04 | < 1e-04 | **** |
| Dorsal | 40 | Middle | Old | 465.000 | 0.569000 | 1.000000 | ns |
| Dorsal | 40 | Middle | Oldest old | 306.000 | < 1e-04 | < 1e-04 | **** |
| Dorsal | 40 | Old | Oldest old | 904.000 | < 1e-04 | < 1e-04 | **** |
| Dorsal | 40 | Young | Middle | 108.000 | 0.326000 | 0.978000 | ns |
| Dorsal | 40 | Young | Old | 393.000 | 0.923000 | 1.000000 | ns |

*Table S4.4: Pairwise Wilcoxon Tests - N1 Amplitude at Each Level by Region (Holm-Bonferroni correction)*

| Region | Level (dB SPL) | Group 1 | Group 2 | W | p | p (adj) | Sig |
|---|---|---|---|---|---|---|---|
| Dorsal | 40 | Young | Oldest old | 318.000 | < 1e-04 | < 1e-04 | **** |
| Dorsal | 50 | Middle | Old | 354.000 | 0.310000 | 0.980000 | ns |
| Dorsal | 50 | Middle | Oldest old | 191.000 | 0.537000 | 1.000000 | ns |
| Dorsal | 50 | Old | Oldest old | 635.000 | 0.080000 | 0.402000 | ns |
| Dorsal | 50 | Young | Middle | 103.000 | 0.245000 | 0.980000 | ns |
| Dorsal | 50 | Young | Old | 276.000 | 0.065000 | 0.388000 | ns |
| Dorsal | 50 | Young | Oldest old | 181.000 | 0.519000 | 1.000000 | ns |
| Dorsal | 60 | Middle | Old | 312.000 | 0.105000 | 0.315000 | ns |
| Dorsal | 60 | Middle | Oldest old | 96.000 | 0.024000 | 0.100000 | ns |
| Dorsal | 60 | Old | Oldest old | 406.000 | 0.224000 | 0.448000 | ns |
| Dorsal | 60 | Young | Middle | 117.000 | 0.510000 | 0.510000 | ns |
| Dorsal | 60 | Young | Old | 244.000 | 0.020000 | 0.100000 | ns |
| Dorsal | 60 | Young | Oldest old | 78.000 | 0.008000 | 0.050000 | ns |
| Dorsal | 70 | Middle | Old | 299.000 | 0.071000 | 0.141000 | ns |
| Dorsal | 70 | Middle | Oldest old | 64.000 | 0.000860 | 0.004000 | ** |
| Dorsal | 70 | Old | Oldest old | 343.000 | 0.042000 | 0.126000 | ns |
| Dorsal | 70 | Young | Middle | 110.000 | 0.363000 | 0.363000 | ns |
| Dorsal | 70 | Young | Old | 230.000 | 0.011000 | 0.045000 | * |
| Dorsal | 70 | Young | Oldest old | 54.000 | 0.000454 | 0.003000 | ** |
| Dorsal | 80 | Middle | Old | 268.000 | 0.024000 | 0.048000 | * |
| Dorsal | 80 | Middle | Oldest old | 37.000 | < 1e-04 | < 1e-04 | **** |
| Dorsal | 80 | Old | Oldest old | 258.000 | 0.002000 | 0.007000 | ** |
| Dorsal | 80 | Young | Middle | 109.000 | 0.345000 | 0.345000 | ns |

*Table S4.4: Pairwise Wilcoxon Tests - N1 Amplitude at Each Level by Region (Holm-Bonferroni correction)*

| Region | Level (dB SPL) | Group 1 | Group 2 | W | p | p (adj) | Sig |
|---|---|---|---|---|---|---|---|
| Dorsal | 80 | Young | Old | 215.000 | 0.006000 | 0.017000 | * |
| Dorsal | 80 | Young | Oldest old | 41.000 | < 1e-04 | 0.000297 | *** |

### Figure 4: Time-Frequency Analysis (30-100 Hz Power)

#### Figure 4D: Peak 30-100 Hz Power Analysis

#### Two-way Repeated Measures ANOVA: Age × Level for 30-100 Hz Power

*Table S5.1: Two-Way Repeated Measures ANOVA - Age × Level for Peak 30-100 Hz Power*

| Region | Stratum | Factor | df1 | df2 | F | p-value | Sig | F (manuscript format) |
|---|---|---|---|---|---|---|---|---|
| Ventral | Between-subjects | Age | 3.000 | 154.000 | 8.248 | <0.001 | *** | F(3, 154) = 8.248 |
| Ventral | Within-subjects | Level | 6.000 | 924.000 | 262.214 | <0.001 | *** | F(6, 924) = 262.214 |
| Ventral | Within-subjects | Age × Level | 18.000 | 924.000 | 7.988 | <0.001 | *** | F(18, 924) = 7.988 |
| Medial | Between-subjects | Age | 3.000 | 230.000 | 5.637 | <0.001 | *** | F(3, 230) = 5.637 |
| Medial | Within-subjects | Level | 6.000 | 1380.000 | 441.000 | <0.001 | *** | F(6, 1380) = 441.000 |
| Medial | Within-subjects | Age × Level | 18.000 | 1380.000 | 31.658 | <0.001 | *** | F(18, 1380) = 31.658 |

| Region | Stratum | Factor | df1 | df2 | F | p-value | Sig | F (manuscript format) |
|---|---|---|---|---|---|---|---|---|
| Dorsal | Between-subjects | Age | 3.000 | 99.000 | 1.273 | 0.288 | ns | F(3, 99) = 1.273 |
| Dorsal | Within-subjects | Level | 6.000 | 594.000 | 143.674 | <0.001 | *** | F(6, 594) = 143.674 |
| Dorsal | Within-subjects | Age × Level | 18.000 | 594.000 | 16.266 | <0.001 | *** | F(18, 594) = 16.266 |

#### Descriptive Statistics: 30-100 Hz Power by Level and Region

*Table S5.2: Descriptive Statistics - Peak 30-100 Hz Power ($\mu V^2/Hz$) by Level, Region, and Age*

| Region | Level (dB SPL) | Age Group | n | Mean | SD | SEM |
|---|---|---|---|---|---|---|
| Ventral | 20 | Young | 25 | 37.72 | 27.05 | 5.41 |
| Ventral | 20 | Middle | 26 | 32.54 | 30.02 | 5.89 |
| Ventral | 20 | Old | 82 | 17.27 | 15.06 | 1.66 |
| Ventral | 20 | Oldest old | 25 | 14.50 | 9.58 | 1.92 |
| Ventral | 30 | Young | 25 | 186.25 | 61.50 | 12.30 |
| Ventral | 30 | Middle | 26 | 124.52 | 120.99 | 23.73 |
| Ventral | 30 | Old | 82 | 37.88 | 45.13 | 4.98 |
| Ventral | 30 | Oldest old | 25 | 18.32 | 8.86 | 1.77 |
| Ventral | 40 | Young | 25 | 260.92 | 111.94 | 22.39 |
| Ventral | 40 | Middle | 26 | 235.55 | 156.96 | 30.78 |
| Ventral | 40 | Old | 82 | 98.23 | 72.89 | 8.05 |
| Ventral | 40 | Oldest old | 25 | 21.51 | 14.45 | 2.89 |
| Ventral | 50 | Young | 25 | 287.15 | 130.99 | 26.20 |
| Ventral | 50 | Middle | 26 | 261.30 | 148.88 | 29.20 |
| Ventral | 50 | Old | 82 | 197.88 | 133.69 | 14.76 |
| Ventral | 50 | Oldest old | 25 | 122.46 | 150.10 | 30.02 |

| Region | Level (dB SPL) | Age Group | n | Mean | SD | SEM |
|---|---|---|---|---|---|---|
| Ventral | 60 | Young | 25 | 301.69 | 139.28 | 27.86 |
| Ventral | 60 | Middle | 26 | 257.51 | 126.53 | 24.82 |
| Ventral | 60 | Old | 82 | 249.20 | 143.20 | 15.81 |
| Ventral | 60 | Oldest old | 25 | 217.23 | 180.57 | 36.11 |
| Ventral | 70 | Young | 25 | 313.88 | 134.02 | 26.80 |
| Ventral | 70 | Middle | 26 | 271.77 | 113.82 | 22.32 |
| Ventral | 70 | Old | 82 | 262.86 | 146.09 | 16.13 |
| Ventral | 70 | Oldest old | 25 | 286.10 | 186.92 | 37.38 |
| Ventral | 80 | Young | 25 | 336.28 | 145.71 | 29.14 |
| Ventral | 80 | Middle | 26 | 274.66 | 104.44 | 20.48 |
| Ventral | 80 | Old | 82 | 269.09 | 137.94 | 15.23 |
| Ventral | 80 | Oldest old | 25 | 296.91 | 175.77 | 35.15 |
| Medial | 20 | Young | 41 | 75.15 | 61.55 | 9.61 |
| Medial | 20 | Middle | 42 | 65.57 | 78.22 | 12.07 |
| Medial | 20 | Old | 111 | 19.93 | 18.68 | 1.77 |
| Medial | 20 | Oldest old | 40 | 11.85 | 5.09 | 0.81 |
| Medial | 30 | Young | 41 | 231.61 | 94.13 | 14.70 |
| Medial | 30 | Middle | 42 | 201.61 | 124.67 | 19.24 |
| Medial | 30 | Old | 111 | 72.30 | 94.43 | 8.96 |
| Medial | 30 | Oldest old | 40 | 13.71 | 4.41 | 0.70 |
| Medial | 40 | Young | 41 | 295.28 | 117.09 | 18.29 |
| Medial | 40 | Middle | 42 | 314.48 | 114.76 | 17.71 |
| Medial | 40 | Old | 111 | 185.03 | 163.28 | 15.50 |
| Medial | 40 | Oldest old | 40 | 24.46 | 16.11 | 2.55 |
| Medial | 50 | Young | 41 | 285.41 | 124.28 | 19.41 |

| Region | Level (dB SPL) | Age Group | n | Mean | SD | SEM |
|---|---|---|---|---|---|---|
| Medial | 50 | Middle | 42 | 310.73 | 108.18 | 16.69 |
| Medial | 50 | Old | 111 | 285.36 | 161.32 | 15.31 |
| Medial | 50 | Oldest old | 40 | 142.43 | 163.76 | 25.89 |
| Medial | 60 | Young | 41 | 293.82 | 132.38 | 20.67 |
| Medial | 60 | Middle | 42 | 299.55 | 107.58 | 16.60 |
| Medial | 60 | Old | 111 | 333.92 | 150.14 | 14.25 |
| Medial | 60 | Oldest old | 40 | 288.28 | 167.05 | 26.41 |
| Medial | 70 | Young | 41 | 295.47 | 137.77 | 21.52 |
| Medial | 70 | Middle | 42 | 299.00 | 104.15 | 16.07 |
| Medial | 70 | Old | 111 | 332.13 | 142.95 | 13.57 |
| Medial | 70 | Oldest old | 40 | 370.47 | 162.16 | 25.64 |
| Medial | 80 | Young | 41 | 300.85 | 144.14 | 22.51 |
| Medial | 80 | Middle | 42 | 294.13 | 103.30 | 15.94 |
| Medial | 80 | Old | 111 | 327.51 | 130.11 | 12.35 |
| Medial | 80 | Oldest old | 40 | 399.13 | 156.23 | 24.70 |
| Dorsal | 20 | Young | 16 | 63.27 | 34.26 | 8.57 |
| Dorsal | 20 | Middle | 17 | 52.28 | 54.61 | 13.24 |
| Dorsal | 20 | Old | 50 | 17.63 | 12.28 | 1.74 |
| Dorsal | 20 | Oldest old | 20 | 10.49 | 5.99 | 1.34 |
| Dorsal | 30 | Young | 16 | 131.34 | 48.20 | 12.05 |
| Dorsal | 30 | Middle | 17 | 123.98 | 74.59 | 18.09 |
| Dorsal | 30 | Old | 50 | 76.16 | 69.93 | 9.89 |
| Dorsal | 30 | Oldest old | 20 | 12.82 | 5.15 | 1.15 |
| Dorsal | 40 | Young | 16 | 147.56 | 60.78 | 15.19 |
| Dorsal | 40 | Middle | 17 | 172.34 | 71.31 | 17.29 |

| Region | Level (dB SPL) | Age Group | n | Mean | SD | SEM |
|---|---|---|---|---|---|---|
| Dorsal | 40 | Old | 50 | 168.39 | 118.99 | 16.83 |
| Dorsal | 40 | Oldest old | 20 | 24.68 | 17.18 | 3.84 |
| Dorsal | 50 | Young | 16 | 146.65 | 71.73 | 17.93 |
| Dorsal | 50 | Middle | 17 | 178.67 | 77.11 | 18.70 |
| Dorsal | 50 | Old | 50 | 225.93 | 140.78 | 19.91 |
| Dorsal | 50 | Oldest old | 20 | 182.07 | 177.99 | 39.80 |
| Dorsal | 60 | Young | 16 | 160.90 | 98.04 | 24.51 |
| Dorsal | 60 | Middle | 17 | 178.07 | 75.15 | 18.23 |
| Dorsal | 60 | Old | 50 | 255.46 | 156.53 | 22.14 |
| Dorsal | 60 | Oldest old | 20 | 315.88 | 192.46 | 43.04 |
| Dorsal | 70 | Young | 16 | 170.58 | 119.96 | 29.99 |
| Dorsal | 70 | Middle | 17 | 179.98 | 77.86 | 18.88 |
| Dorsal | 70 | Old | 50 | 262.24 | 150.76 | 21.32 |
| Dorsal | 70 | Oldest old | 20 | 350.71 | 168.29 | 37.63 |
| Dorsal | 80 | Young | 16 | 169.74 | 127.81 | 31.95 |
| Dorsal | 80 | Middle | 17 | 175.41 | 82.99 | 20.13 |
| Dorsal | 80 | Old | 50 | 265.42 | 143.72 | 20.32 |
| Dorsal | 80 | Oldest old | 20 | 397.46 | 154.10 | 34.46 |

#### Post-hoc Tests: Age Effect at Each Level by Region (30-100 Hz)

*Table S5.3: Kruskal-Wallis Tests - Age Effect on 30-100 Hz Power at Each Level by Region*

| Region | Level (dB SPL) | H | df | p-value | Sig |
|---|---|---|---|---|---|
| Ventral | 20 | 29.432 | 3 | <0.001 | *** |
| Ventral | 30 | 84.290 | 3 | <0.001 | *** |
| Ventral | 40 | 87.816 | 3 | <0.001 | *** |

| Region | Level (dB SPL) | H | df | p-value | Sig |
|---|---|---|---|---|---|
| Ventral | 50 | 28.954 | 3 | <0.001 | *** |
| Ventral | 60 | 6.948 | 3 | 0.0736 | ns |
| Ventral | 70 | 3.011 | 3 | 0.39 | ns |
| Ventral | 80 | 4.040 | 3 | 0.257 | ns |
| Medial | 20 | 72.360 | 3 | <0.001 | *** |
| Medial | 30 | 124.677 | 3 | <0.001 | *** |
| Medial | 40 | 111.476 | 3 | <0.001 | *** |
| Medial | 50 | 40.089 | 3 | <0.001 | *** |
| Medial | 60 | 5.086 | 3 | 0.166 | ns |
| Medial | 70 | 7.412 | 3 | 0.0599 | ns |
| Medial | 80 | 14.251 | 3 | 0.00258 | ** |
| Dorsal | 20 | 27.912 | 3 | <0.001 | *** |
| Dorsal | 30 | 47.555 | 3 | <0.001 | *** |
| Dorsal | 40 | 39.189 | 3 | <0.001 | *** |
| Dorsal | 50 | 6.022 | 3 | 0.111 | ns |
| Dorsal | 60 | 10.549 | 3 | 0.0144 | * |
| Dorsal | 70 | 17.208 | 3 | <0.001 | *** |
| Dorsal | 80 | 26.017 | 3 | <0.001 | *** |

#### Pairwise Comparisons: 30-100 Hz Power at Each Level by Region

*Table S5.4: Pairwise Wilcoxon Tests – 30-100 Hz Power at Each Level by Region (Holm-Bonferroni correction)*

| Region | Level (dB SPL) | Group 1 | Group 2 | W | p | p (adj) | Sig |
|---|---|---|---|---|---|---|---|
| Ventral | 20 | Young | Middle | 426.0000 | 0.0580 | 0.2310 | ns |
| Ventral | 20 | Young | Old | 1735.0000 | 0.0000 | 0.0000 | **** |
| Ventral | 20 | Young | Oldest old | 564.0000 | 0.0000 | 0.0000 | **** |

| Region | Level (dB SPL) | Group 1 | Group 2 | W | p | p (adj) | Sig |
|---|---|---|---|---|---|---|---|
| Ventral | 20 | Middle | Old | 1258.0000 | 0.1690 | 0.5010 | ns |
| Ventral | 20 | Middle | Oldest old | 399.0000 | 0.1670 | 0.5010 | ns |
| Ventral | 20 | Old | Oldest old | 1146.0000 | 0.3750 | 0.5010 | ns |
| Ventral | 30 | Young | Middle | 497.0000 | 0.0009 | 0.0020 | ** |
| Ventral | 30 | Young | Old | 1982.0000 | 0.0000 | 0.0000 | **** |
| Ventral | 30 | Young | Oldest old | 624.0000 | 0.0000 | 0.0000 | **** |
| Ventral | 30 | Middle | Old | 1839.0000 | 0.0000 | 0.0000 | **** |
| Ventral | 30 | Middle | Oldest old | 637.0000 | 0.0000 | 0.0000 | **** |
| Ventral | 30 | Old | Oldest old | 1381.0000 | 0.0090 | 0.0090 | ** |
| Ventral | 40 | Young | Middle | 406.0000 | 0.1300 | 0.1300 | ns |
| Ventral | 40 | Young | Old | 1840.0000 | 0.0000 | 0.0000 | **** |
| Ventral | 40 | Young | Oldest old | 624.0000 | 0.0000 | 0.0000 | **** |
| Ventral | 40 | Middle | Old | 1767.0000 | 0.0000 | 0.0000 | **** |
| Ventral | 40 | Middle | Oldest old | 649.0000 | 0.0000 | 0.0000 | **** |
| Ventral | 40 | Old | Oldest old | 1851.0000 | 0.0000 | 0.0000 | **** |
| Ventral | 50 | Young | Middle | 385.0000 | 0.2650 | 0.2650 | ns |
| Ventral | 50 | Young | Old | 1473.0000 | 0.0010 | 0.0030 | ** |
| Ventral | 50 | Young | Oldest old | 510.0000 | 0.0001 | 0.0004 | *** |
| Ventral | 50 | Middle | Old | 1369.0000 | 0.0300 | 0.0590 | ns |
| Ventral | 50 | Middle | Oldest old | 521.0000 | 0.0001 | 0.0007 | *** |
| Ventral | 50 | Old | Oldest old | 1534.0000 | 0.0002 | 0.0007 | *** |
| Ventral | 60 | Young | Middle | 388.0000 | 0.2410 | 0.4820 | ns |
| Ventral | 60 | Young | Old | 1281.0000 | 0.0600 | 0.3000 | ns |
| Ventral | 60 | Young | Oldest old | 427.0000 | 0.0260 | 0.1570 | ns |
| Ventral | 60 | Middle | Old | 1137.0000 | 0.6120 | 0.6120 | ns |

| Region | Level (dB SPL) | Group 1 | Group 2 | W | p | p (adj) | Sig |
|---|---|---|---|---|---|---|---|
| Ventral | 60 | Middle | Oldest old | 404.0000 | 0.1400 | 0.4760 | ns |
| Ventral | 60 | Old | Oldest old | 1237.0000 | 0.1190 | 0.4760 | ns |
| Ventral | 70 | Young | Middle | 380.0000 | 0.3070 | 1.0000 | ns |
| Ventral | 70 | Young | Old | 1275.0000 | 0.0660 | 0.3970 | ns |
| Ventral | 70 | Young | Oldest old | 338.0000 | 0.6310 | 1.0000 | ns |
| Ventral | 70 | Middle | Old | 1165.0000 | 0.4790 | 1.0000 | ns |
| Ventral | 70 | Middle | Oldest old | 323.0000 | 0.9780 | 1.0000 | ns |
| Ventral | 70 | Old | Oldest old | 984.0000 | 0.7660 | 1.0000 | ns |
| Ventral | 80 | Young | Middle | 399.0000 | 0.1670 | 0.8350 | ns |
| Ventral | 80 | Young | Old | 1307.0000 | 0.0380 | 0.2290 | ns |
| Ventral | 80 | Young | Oldest old | 330.0000 | 0.7440 | 1.0000 | ns |
| Ventral | 80 | Middle | Old | 1149.0000 | 0.5530 | 1.0000 | ns |
| Ventral | 80 | Middle | Oldest old | 300.0000 | 0.6470 | 1.0000 | ns |
| Ventral | 80 | Old | Oldest old | 937.0000 | 0.5190 | 1.0000 | ns |
| Medial | 20 | Young | Middle | 1146.0000 | 0.0090 | 0.0180 | * |
| Medial | 20 | Young | Old | 4035.0000 | 0.0000 | 0.0000 | **** |
| Medial | 20 | Young | Oldest old | 1593.0000 | 0.0000 | 0.0000 | **** |
| Medial | 20 | Middle | Old | 3141.0000 | 0.0009 | 0.0030 | ** |
| Medial | 20 | Middle | Oldest old | 1263.0000 | 0.0001 | 0.0002 | *** |
| Medial | 20 | Old | Oldest old | 2838.0000 | 0.0090 | 0.0180 | * |
| Medial | 30 | Young | Middle | 1016.0000 | 0.1600 | 0.1600 | ns |
| Medial | 30 | Young | Old | 4070.0000 | 0.0000 | 0.0000 | **** |
| Medial | 30 | Young | Oldest old | 1640.0000 | 0.0000 | 0.0000 | **** |
| Medial | 30 | Middle | Old | 3926.0000 | 0.0000 | 0.0000 | **** |
| Medial | 30 | Middle | Oldest old | 1680.0000 | 0.0000 | 0.0000 | **** |

| Region | Level (dB SPL) | Group 1 | Group 2 | W | p | p (adj) | Sig |
|---|---|---|---|---|---|---|---|
| Medial | 30 | Old | Oldest old | 3539.0000 | 0.0000 | 0.0000 | **** |
| Medial | 40 | Young | Middle | 772.0000 | 0.4220 | 0.4220 | ns |
| Medial | 40 | Young | Old | 3424.0000 | 0.0000 | 0.0000 | **** |
| Medial | 40 | Young | Oldest old | 1640.0000 | 0.0000 | 0.0000 | **** |
| Medial | 40 | Middle | Old | 3619.0000 | 0.0000 | 0.0000 | **** |
| Medial | 40 | Middle | Oldest old | 1680.0000 | 0.0000 | 0.0000 | **** |
| Medial | 40 | Old | Oldest old | 4049.0000 | 0.0000 | 0.0000 | **** |
| Medial | 50 | Young | Middle | 753.0000 | 0.3290 | 0.6580 | ns |
| Medial | 50 | Young | Old | 2352.0000 | 0.7520 | 0.7520 | ns |
| Medial | 50 | Young | Oldest old | 1344.0000 | 0.0000 | 0.0000 | **** |
| Medial | 50 | Middle | Old | 2687.0000 | 0.1460 | 0.4380 | ns |
| Medial | 50 | Middle | Oldest old | 1425.0000 | 0.0000 | 0.0000 | **** |
| Medial | 50 | Old | Oldest old | 3518.0000 | 0.0000 | 0.0000 | **** |
| Medial | 60 | Young | Middle | 833.0000 | 0.8030 | 1.0000 | ns |
| Medial | 60 | Young | Old | 1925.0000 | 0.1460 | 0.7300 | ns |
| Medial | 60 | Young | Oldest old | 884.0000 | 0.5510 | 1.0000 | ns |
| Medial | 60 | Middle | Old | 2018.0000 | 0.2010 | 0.8040 | ns |
| Medial | 60 | Middle | Oldest old | 937.0000 | 0.3730 | 1.0000 | ns |
| Medial | 60 | Old | Oldest old | 2661.0000 | 0.0630 | 0.3790 | ns |
| Medial | 70 | Young | Middle | 821.0000 | 0.7200 | 0.7200 | ns |
| Medial | 70 | Young | Old | 1913.0000 | 0.1330 | 0.5320 | ns |
| Medial | 70 | Young | Oldest old | 587.0000 | 0.0280 | 0.1430 | ns |
| Medial | 70 | Middle | Old | 1984.0000 | 0.1570 | 0.5320 | ns |
| Medial | 70 | Middle | Oldest old | 597.0000 | 0.0240 | 0.1430 | ns |
| Medial | 70 | Old | Oldest old | 1909.0000 | 0.1900 | 0.5320 | ns |

| Region | Level (dB SPL) | Group 1 | Group 2 | W | p | p (adj) | Sig |
|---|---|---|---|---|---|---|---|
| Medial | 80 | Young | Middle | 838.0000 | 0.8380 | 0.8380 | ns |
| Medial | 80 | Young | Old | 1948.0000 | 0.1750 | 0.3960 | ns |
| Medial | 80 | Young | Oldest old | 534.0000 | 0.0070 | 0.0290 | * |
| Medial | 80 | Middle | Old | 1962.0000 | 0.1320 | 0.3960 | ns |
| Medial | 80 | Middle | Oldest old | 467.0000 | 0.0004 | 0.0030 | ** |
| Medial | 80 | Old | Oldest old | 1566.0000 | 0.0060 | 0.0290 | * |
| Dorsal | 20 | Young | Middle | 170.0000 | 0.2310 | 0.3040 | ns |
| Dorsal | 20 | Young | Old | 715.0000 | 0.0000 | 0.0000 | **** |
| Dorsal | 20 | Young | Oldest old | 301.0000 | 0.0000 | 0.0000 | **** |
| Dorsal | 20 | Middle | Old | 525.0000 | 0.1520 | 0.3040 | ns |
| Dorsal | 20 | Middle | Oldest old | 244.0000 | 0.0240 | 0.0930 | ns |
| Dorsal | 20 | Old | Oldest old | 675.0000 | 0.0230 | 0.0930 | ns |
| Dorsal | 30 | Young | Middle | 145.0000 | 0.7630 | 0.7630 | ns |
| Dorsal | 30 | Young | Old | 616.0000 | 0.0010 | 0.0040 | ** |
| Dorsal | 30 | Young | Oldest old | 320.0000 | 0.0000 | 0.0000 | **** |
| Dorsal | 30 | Middle | Old | 603.0000 | 0.0110 | 0.0210 | * |
| Dorsal | 30 | Middle | Oldest old | 340.0000 | 0.0000 | 0.0000 | **** |
| Dorsal | 30 | Old | Oldest old | 897.0000 | 0.0000 | 0.0000 | **** |
| Dorsal | 40 | Young | Middle | 104.0000 | 0.2600 | 0.7800 | ns |
| Dorsal | 40 | Young | Old | 390.0000 | 0.8870 | 0.9340 | ns |
| Dorsal | 40 | Young | Oldest old | 318.0000 | 0.0000 | 0.0000 | **** |
| Dorsal | 40 | Middle | Old | 476.0000 | 0.4670 | 0.9340 | ns |
| Dorsal | 40 | Middle | Oldest old | 332.0000 | 0.0000 | 0.0000 | **** |
| Dorsal | 40 | Old | Oldest old | 924.0000 | 0.0000 | 0.0000 | **** |
| Dorsal | 50 | Young | Middle | 97.0000 | 0.1680 | 0.6720 | ns |

| Region | Level (dB SPL) | Group 1 | Group 2 | W | p | p (adj) | Sig |
|---|---|---|---|---|---|---|---|
| Dorsal | 50 | Young | Old | 272.0000 | 0.0560 | 0.3380 | ns |
| Dorsal | 50 | Young | Oldest old | 182.0000 | 0.4980 | 0.9330 | ns |
| Dorsal | 50 | Middle | Old | 364.0000 | 0.3830 | 0.9330 | ns |
| Dorsal | 50 | Middle | Oldest old | 204.0000 | 0.3110 | 0.9330 | ns |
| Dorsal | 50 | Old | Oldest old | 639.0000 | 0.0720 | 0.3590 | ns |
| Dorsal | 60 | Young | Middle | 109.0000 | 0.3450 | 0.4480 | ns |
| Dorsal | 60 | Young | Old | 241.0000 | 0.0180 | 0.0880 | ns |
| Dorsal | 60 | Young | Oldest old | 78.0000 | 0.0080 | 0.0500 | * |
| Dorsal | 60 | Middle | Old | 322.0000 | 0.1400 | 0.4200 | ns |
| Dorsal | 60 | Middle | Oldest old | 98.0000 | 0.0280 | 0.1120 | ns |
| Dorsal | 60 | Old | Oldest old | 406.0000 | 0.2240 | 0.4480 | ns |
| Dorsal | 70 | Young | Middle | 103.0000 | 0.2450 | 0.2450 | ns |
| Dorsal | 70 | Young | Old | 229.0000 | 0.0110 | 0.0430 | * |
| Dorsal | 70 | Young | Oldest old | 54.0000 | 0.0005 | 0.0030 | ** |
| Dorsal | 70 | Middle | Old | 303.0000 | 0.0800 | 0.1600 | ns |
| Dorsal | 70 | Middle | Oldest old | 65.0000 | 0.0010 | 0.0050 | ** |
| Dorsal | 70 | Old | Oldest old | 343.0000 | 0.0420 | 0.1260 | ns |
| Dorsal | 80 | Young | Middle | 107.0000 | 0.3090 | 0.3090 | ns |
| Dorsal | 80 | Young | Old | 216.0000 | 0.0060 | 0.0180 | * |
| Dorsal | 80 | Young | Oldest old | 41.0000 | 0.0001 | 0.0003 | *** |
| Dorsal | 80 | Middle | Old | 270.0000 | 0.0260 | 0.0520 | ns |
| Dorsal | 80 | Middle | Oldest old | 37.0000 | 0.0000 | 0.0001 | **** |
| Dorsal | 80 | Old | Oldest old | 258.0000 | 0.0020 | 0.0070 | ** |
